# R6G narrows BmrA conformational spectrum for a more efficient use of ATP

**DOI:** 10.1101/2024.03.15.585201

**Authors:** A Gobet, L Moissonnier, E Zarkadas, S Magnard, E Bettler, J Martin, R Terreux, G Schoehn, C Orelle, JM Jault, P Falson, V Chaptal

## Abstract

Multidrug ABC transporters harness the energy of ATP binding and hydrolysis to change conformation and thereby translocate substrates out of the cell to detoxify them. While this general access mechanism scheme is well accepted, molecular details of this interplay is still elusive. Rhodamine6G binding on a catalytic mutant of the homodimeric multidrug ABC transporter BmrA triggers a cooperative binding of ATP on the two identical nucleotide-binding-sites, otherwise Michaelian. We investigated this asymmetric behavior via a structural-enzymology approach, solving cryoEM structure of BmrA at defined ATP ratio along the enzymatic transition, highlighting the plasticity of BmrA as it undergoes the transition from inward to outward facing conformations. Analysis of continuous heterogeneity within cryoEM data and structural dynamics, revealed that Rhodamine6G narrows the conformational spectrum explored by the nucleotide-binding-domains, describing the allosteric effect of drug binding that optimizes the ATP-dependent conversion of the transporter to the outward-facing state. Following on these findings, the effect of drug-binding showed an ATPase stimulation and a maximal transport activity of the wild-type protein at the concentration-range where the allosteric transition occurs. Drug diffusion rate is the likely rate-limiting step of the reaction, while drug transport and ATPase activities are in effect uncoupled.

## 1. Introduction

Drug resistance mediated by ABC (ATP-Binding Cassette) transporters contributes to the first line of defense for organisms, decreasing the intracellular drug concentration and allowing cells to further adapt by acquiring target mutations [1–3]. A typical hallmark of multidrug ABC transporters is the ability to recognize and transport a wide array of structurally unrelated substrates across the plasma membrane, thereby protecting the organism against many xenobiotics. These transporters thus harbor a polyspecificity for multiple ligands, that they can accommodate within a rather large and size adaptable binding pocket. The wealth of structural data available on these transporters as well as the biochemical and biophysical characterizations have highlighted their plasticity and their ability to adapt to various ligands [4–9]. At the core, it is clear that ABC transporters are very flexible and undergo significant conformational changes to perform their transport function.

Type IV ABC transporters involved in MultiDrug Resistance (MDR) phenotype perform their function by undergoing a series of conformational changes pertaining to the alternating access mechanism. They are formed of two trans-membrane domains (TMD) forming a cavity at their center where substrates bind, and of two nucleotide-binding domains (NBD) binding and hydrolyzing ATP-Mg^2+^ to fuel the system. At resting state, they populate an Inward-Facing (IF) conformation where substrates can bind. Binding of ATP-Mg^2+^ to the NBDs bring them together, translating this conformational change to the TMDs which then open towards the outside (Outward-Facing, OF) where substrates are released. ATP hydrolysis and product release reset the transporter back to the IF conformation, ready for another cycle. While the overall scheme of this transport cycle is widely accepted, details of each step and their order are still highly debated.

The *Bacillus subtilis* efflux pump BmrA, belongs to the type-IV family of MDR ABC transporters[10, 11], conferring resistance to cervimycin-C secreted by the biotope competitor *Streptomyces tendæ*[12]. A *B. subtilis* strain resistant to the antibiotic was isolated and contained mutations on the *bmra* promotor region increasing and stabilizing its mRNA, which results in BmrA overexpression at the membrane[12, 13]. BmrA binds and transports many structurally-unrelated drugs, including doxorubicin or daunorubicin, Hœchst33342, Rhodamine6G (R6G) or ethidium bromide[11, 13]. BmrA harbors a demonstrated plasticity to handle its ligands, thus conferring an evolutionary advantage to cells. It has been postulated that instead of going through discrete well-defined states, it uses its intrinsic plasticity to swing back and forth around an intermediate occluded state while deforming to adapt to each transported molecule[10, 13–17]. However so far, the molecular details of such critical plasticity and how it couples both functions of drug translocation and ATP hydrolysis lacks detailed structural information.

Here, we get insights on BmrA plasticity undergoing the IF to OF conformational change and its ability to respond to ligand/substrate binding using a structural enzymology approach. We first probed ATP-Mg^2+^ binding to BmrA in the absence or presence of R6G, showing a shift from Michaelian to allosteric behavior. We then visualized by cryoEM the effects of both ligand binding along this transition. We further analyzed protein flexibility within cryoEM data using a software we developed [18], and reproduced these findings by Molecular Dynamics (MD) simulations. Data show that R6G focuses BmrA conformational changes, allowing for a more efficient space exploration. Finally, we could establish that drug binding and shift in flexibility results in ATPase activation for ATP concentration where the transition occurs, resulting in a tighter coupling of transport and ATPase activities, otherwise uncoupled.

## 2. Results

### 2.1 Shift from Michaelian to allosteric ATP-Mg^2+^ binding in response to R6G binding

BmrA is a dimer, each monomer bearing 1 TMD and 1 NBD [19]. The two NBDs are thus identical, and the OF structure of the E504A mutant we previously solved confirms that they bind ATP-Mg^2+^ identically[13], in accordance with all other structures of type-IV ABC transporters. Probing ATP-Mg^2+^ binding (Figure 1A) indeed reveals a Michaelian type of binding (hyperbolic-type curve, K_d-app_ = 154.0 µM ± 49.0). Importantly, the E504A catalytic mutant that still binds ATP but cannot hydrolyze it [20] was used in this study; this thus allowed to probe a unidirectional conformational transistion from IF to OF states, BmrA being blocked when reaching the OF conformation when complexed with ATP-Mg^2+^. Interestingly, addition of the substrate R6G prior to ATP-Mg^2+^ binding resulted in a shift towards a sigmoidal-type curve, suggesting an allosteric binding of ATP-Mg^2+^ (Figure 1C), and meaning that although identical the NBDs do not bind ATP-Mg^2+^ the same way anymore (K_0.5_ = 70.0 µM ± 2.6, n=3.4 ± 0.3). In addition, the pre-binding of R6G increased the apparent affinity for ATP, together with the steep transition suggesting that R6G binding ensures a more efficient conversion to the OF conformation mediated by ATP-Mg^2+^ binding. Since this behavior has consequences on the transport cycle, we performed a structural enzymology study to investigate this transition structurally. CryoEM grids were frozen at key ligand concentrations and imaged; without ATP-Mg^2+^ in absence or presence of R6G (named E504A^apo^ and E504A^R6G^), at a 1:1 molar ratio ATP-Mg^2+^:BmrA dimer (E504A^apo-25µMATP^ and E504A^R6G-25µMATP^), at the K_0.5_ or close to the K_d-app_ for ATP-Mg^2+^ (E504A^apo-100µMATP^ and E504A^R6G-70µMATP^), and at saturating concentrations of ATP-Mg^2+^ (E504A^apo-5mMATP^ and E504A^R6G-5mMATP^) (Figure 1, Supp Figures 1-8). Without nucleotide, BmrA is 100% in the IF conformation, in good agreement with SANS and HDX results [16], as well as cryoEM structures of the WT protein and A582C mutant [21] and single molecule FRET recent studies [22]. Progressive addition of ATP-Mg^2+^ shifts the population towards the OF conformation to reach 100% of the population at saturating ATP concentrations. At a 1:1 molar ratio ATP-Mg^2+^:BmrA in the presence of R6G, 25% of BmrA population already shifts towards the OF conformation. This is in good agreement with the allostery model, and the notion that ATP-Mg^2+^ binding to one monomer increases the affinity of ATP-Mg^2+^ for the other monomer (Figure 1D). Two molecules of ATP-Mg^2+^ are thus trapped in the OF conformation, being 50% of the ATP-Mg^2+^ in the sample, and the clear shape of the OF conformation makes it distinguishable in the reconstructions. In contrast, in the absence of R6G (E504A^apo-25µMATP^), only IF reconstructions could be observed as ATP-Mg^2+^ will distribute equally among both NBDs and will not yield enough particles in the OF conformation to be seen (Figure 1B). These results are consistent with the transition curve observed. At the K_0.5_ for ATP-Mg^2+^, 60% of the population is switched to the OF conformation in presence of R6G (E504A^R6G-70µMATP^), while greater concentrations of ATP-Mg^2+^ are required to reach a similar population distribution in the absence of drug. Altogether, this structural enzymology approach exemplifies how R6G binding on BmrA results on a more efficient conversion from IF to OF conformation, on a narrower range of ATP-Mg^2+^ concentration.

**Figure 1:**
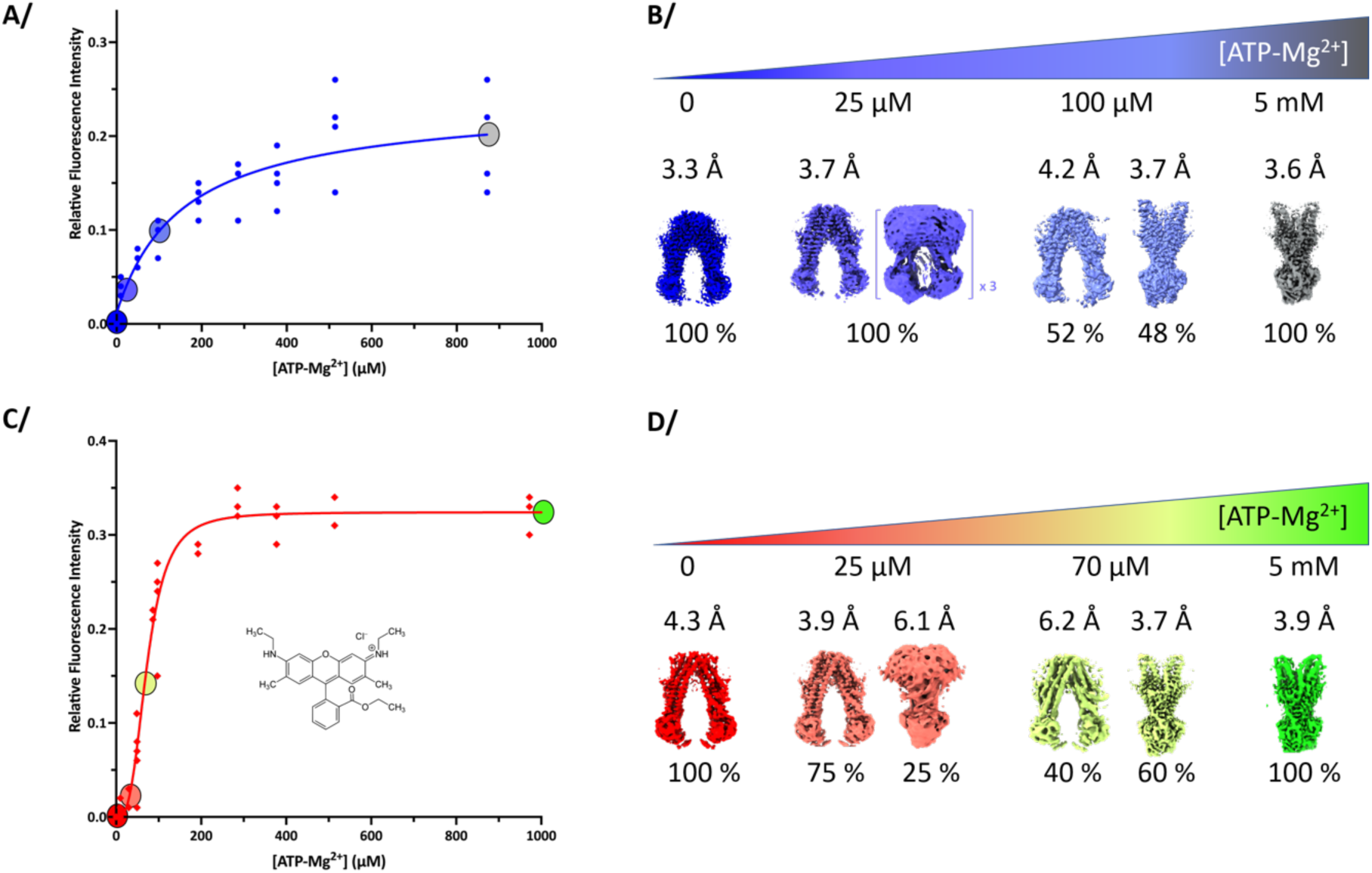
Binding of ATP-Mg^2+^ on BmrA E504A in absence or presence of R6G and corresponding maps. **A/** Binding curves of ATP-Mg^2+^ in absence of Rhodamine 6G is Michaelian. Circles correspond to the conditions at which cryo-EM data were collected. Blue to grey circles correspond to different condition of collected cryo-EM data at 0 µM, 25 µM, 100 µM and 5 mM ATP-Mg^2+^ for 25µM of BmrA dimer in the sample. **B/** Coulomb potential maps corresponding to each condition of collect and their relative proportions. For E504A^apo-25µMATP^, one reconstruction could yield high resolution and 3 additional volumes could be resolved to IF conformations. **C/** Binding curve of ATP-Mg^2+^ in presence of 100 µM R6G is allosteric. Cryo-EM data were collected at different concentration of ATP-Mg^2+^. Red to green circles correspond to the four conditions: 0 µM, 25 µM, 70 µM and 5 mM ATP-Mg^2+^. **D/** Coulomb potential maps corresponding to each condition and their relative proportions.

### 2.2 Flexibility of BmrA in the IF and OF conformations

3D models were built into each high-resolution 3D reconstruction to gain molecular insights on the IF to OF conversion (Figure 2, Supp-Figures 9). Clear electron densities could be observed for R6G in all the IF and OF reconstructions where it was added. In contrast, all reconstructions where R6G was absent did not show density in the substrate-binding cavity (Supp-Figure 10). For the OF conformation, current observations represent a biological duplicate of observations we previously made in [13], with R6G wedged between TM1-2 and TM5’-6’ of each half transporter, with a total of 2 R6G per BmrA dimer (Figure 2DE). Binding of R6G triggers a closure of TM1 towards TM2, reinforcing the hand-fan motion of these 2 TMs previously hypothesized [13]. Admittedly, density for R6G is not very well defined, allowing for multiple positioning of the molecule, which denotes intrinsic R6G flexibility within the drug-binding pocket, as was also observed by molecular dynamics (MD) simulations [13]. In the IF conformations, R6G could also be modelled within the TMD in-between the same TM helices. Here again, electron density denotes multiple positions adopted by the molecule (also observed in MD simulations below), in line with the ability of BmrA to recognize structurally-unrelated ligands. To exemplify this, we have modelled R6G in 2 conformations of the benzoate moiety with 180° flip of the xanthene core fitting equally well (Figure 2C, left). In E504A^apo^, E504A^apo-25µMATP^ and E504A^apo-100µMATP^, multiple IF conformations of BmrA are observed with a rearrangement of TM helices and the NBD (Figure 2A-C), in line with the flexibility observed for BmrA [16, 21]and other ABC transporters in this conformation [4, 7, 23]. Notably, the E504A^apo^ structure is exactly the same as the WT or A582C mutant structures recently solved (rmsd = 0.52Å over 902 atoms and 0.45 Å over 883 atoms, respectively; suppFigure 11 [21]). R6G binding in E504A^R6G^ and E504A^R6G-25µMATP^ results in a further rearrangement of TM helices within the dimer, with a wider opening of TM4-5-6’. Surprisingly, the rearrangement does not occur within a monomer, as it overlays well with BmrA in absence of R6G (Figure 2C, right part of the structure on which the overlay is performed). The effect of R6G binding further translates all the way to the NBD, resulting in a reorientation of the 2 NBD that face each other more directly (Figure 2B), hinting at an influence on ATP binding observed in Figure 1. Altogether, these structures highlight the flexibility of BmrA in the IF conformation, and the fact that R6G has an allosteric effect that reorients the protein at the ATP-binding site level, reminiscent of what was observed for ABCB1 in presence of drug or inhibitor [24].

**Figure 2:**
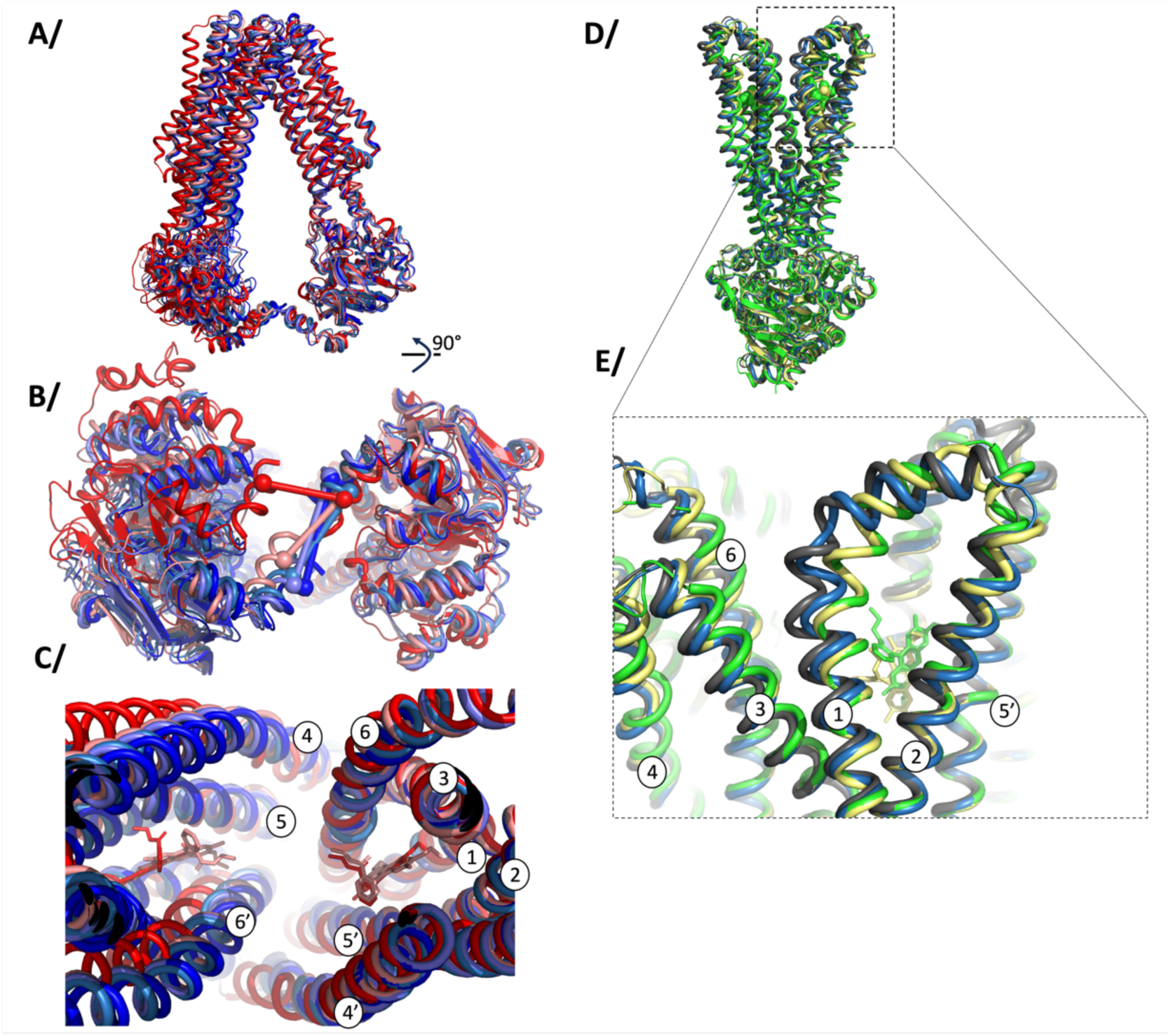
Superposition of models in IF or OF conformations. Colors match the ones of Figure 1 and Supp-Fig 9. Protein in cartoon and ligands are in sticks, TM helices are numbered when appropriate. **A/**, **B**/ and **C/** BmrA in the IF conformation seen from the side, from under the NBD or a close up of the substrate-binding pocket, respectively. Overlay was performed on the 3 first TM helices of chain A, residues 1-161. On panel B/, a line is drawn between residues 577 of each monomer to highlight the conformational change. **D/** and **E/** show BmrA in the OF conformation, with **E/** being a close-up view of the TM1-2 loop with R6G in its pocket. Overlay was performed on the last 4 helices of chain A, residues 104-300 as in [13].

### 2.3 Impact of R6G on BmrA dynamics and conformational space exploration

With cryoEM structure solution, it is possible to visualize inter-particle difference within a reconstruction, allowing to deconvolute protein movements in several directions of latent space[25]. Note that these changes can be observed through different methods available in many software, nicely reviewed in [26]. Importantly, the movements that can be observed result from different interpretation of the latent space and on how to deconvolute the movement. Software using neural networks to deconvolute movements, such as mannifoldEM, 3DFlex, CryoDRGN [27–29] or others use non-linear interpretation of the data to show deformations linking sub-states within the particle stack. These methods are very powerful to highlight special deformations undergone by the protein within each sub-state but will not model the link between sub-states. In a different manner, linear models of conformational heterogeneity, such as multibody refinement or 3DVA [25, 30], perform a principal component analysis of particle variance compared to the consensus reconstruction. This later analysis allows to decode general types of movements present within the particle stack, decomposing complex movements into simplified motions; the global movements undergone by the protein are thus visualized through several main components, allowing for a deconvolution of a complex movement into simplified sub-movements. This later analysis was chosen to analyze the current datasets as it can reveal the overall influence of R6G on BmrA deformations and conformational space exploration. 3DVA was computed in simple mode, resulting in adding weights to the consensus map in latent space directions calculated by principal component analysis, or intermediate mode calculating real maps along the same direction. For this case, both methods gave the same result.

Such calculation was applied to E504A^apo^ and E504A^R6G^ to visualize movements within BmrA as a response to R6G binding (Supp-movies). Of note, the particle stacks of these two datasets show similar distribution of particles in 3D (no special orientation of the protein within the ice), and a gaussian distribution in latent space, allowing to compare similar types of particle distribution, making the movements directly comparable (Supp-Figure 12). To fully understand underlying movements, we designed a software that refines an ensemble of models inside this ensemble of maps [31] and applied the procedure to BmrA (Supp-figure 13). Each component shows different movements, with clear rotations and translations of the NBD, that originate from the kinks in TM helices (Figure 3AB, Supp-Figures 14-16). For E504A^apo^, three types of movements could be inferred. First a 12° rotation of the NBD in the continuation the TM helices, centered on the NBD beta sheet (V363), is observed in the first component (component 0). Second, a 16° rotation of the NBD alone centered on E453 is observed, without movement of TM helices (Component 1). Third, an overall oscillation of the structure is observed with translations leading the NBDs 5 Å away from each other (component 2). In presence of R6G (E504A^R6G^) similar protein motions are observed and can thus be directly compared to E504A^apo^; the principal components in which these movements appear are different as expected for this type of analysis. The first NBD rotation centered on V363 is conserved in E504A^R6G^, while the rotation centered on E453 is reduced by more than half. Notably, the oscillations that bring the NBDs closer together is greatly affected as the E504A^R6G^ structure is stiffened and movements are very reduced (less than 1 Å) (Figure 3AB). This analysis shows that 1/ the NBDs explore a large conformational space by undergoing rotations and translations. The presence of R6G influences one rotation by reducing half of its amplitude; importantly the NBD movements start from a different starting point in presence of R6G with the NBDs already oriented closer to one another. 2/ The presence of R6G stabilizes the NBDs slightly further apart with reduced translation in this direction. Overall, R6G decreases the space exploration of the NBDs. At first, this interpretation might seem counter-intuitive as we would expect NBDs to close more easily according to the allostery model, but one has to keep in mind the different time-scale of the experiment. Further investigations below bring more sense to this observation.

We then subjected both proteins to MD simulations in a lipid bilayer to investigate movements in the IF conformation linked to R6G binding (Figure 3C-F, Supp-Figures 17-22). ATP-Mg^2+^ was added in an attempt to induce full closure. All replicates show BmrA movements towards a closure of the drug-binding cavity and the NBDs getting closer to each other. The amplitudes of the movements are in different range in each replicate, but overall, all the distances aggregated show that both E504A^apo^ and E504A^R6G^ are able to move with similar range (Supp-Figure 20); the drug thus does not prevent BmrA overall movements. However, the conformational space explored by E504A^R6G^ is much more focused compared to E504A^apo^ in which NBDs are moving in many directions (Supp-Figures 19), showing the allosteric influence of R6G on the NBDs. Simulations also reveal that R6G is mobile within its binding pocket but only explores the nearby space of its original position, owing to the hydrophobicity of the substrate-binding pocket (Supp-Figure 22). When R6G is present, TM helices mostly stay near their original positions, suggesting that R6G prevents direct closure of helices due to steric hindrance, while a stronger closure is observed in its absence. Altogether, the dynamic studies of R6G effect on NBD closure indicate that it reduces and focuses the conformational space explored by BmrA in the IF conformation.

**Figure 3:**
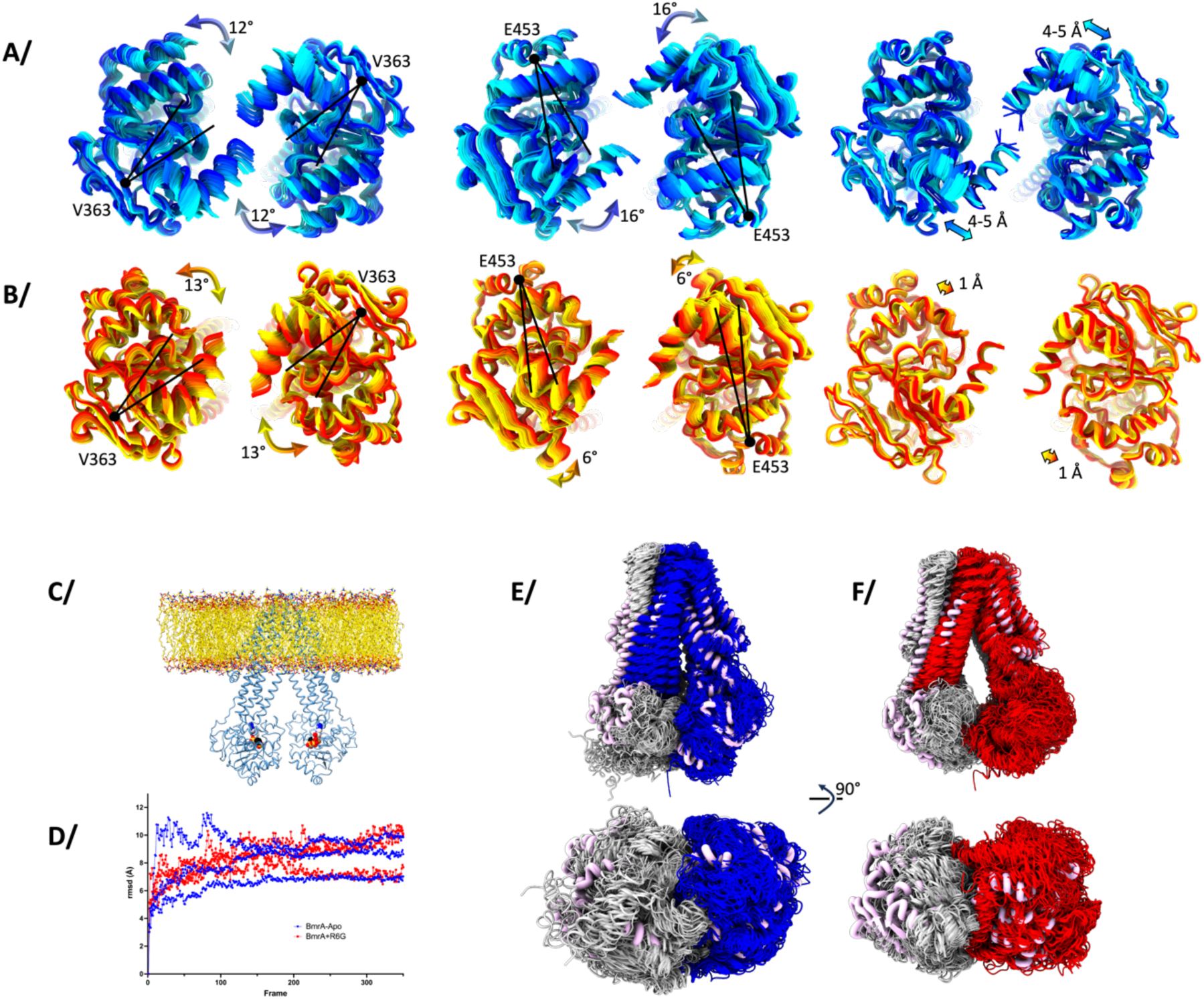
Dynamics of BmrA in the IF conformation. **A/** 3DVA analysis of E504A^apo^ in the first 3 components, resulting in 20 maps each. Models built by variability refinement in the maps are represented in cartoon and colored from blue to cyan, with view from the NBDs. The main movement is represented on the structure with black lines representing the NBD rotation and the colored arrows depict the movement and its amplitude. Details of this analysis are shown in Supp-Figure 17-22. **B/** same as A/ for E504A^R6G^, colored from red to yellow. **C/** BmrA was inserted in a lipid bilayer and ATP-Mg^2+^ was added to start MD simulation, with or without R6G. **D/** Rmsd of the MD trajectories for 3 replicates. The color code corresponds to E/ and F/. **E/** Overlay of 3 replications of MD simulations of BmrA apo to show the conformations sampled during the simulations. Thick pink ribbons correspond to the initial structure. **F/** same as E/ for BmrA in presence of R6G.

### 2.4 Impact of R6G on ATPase and transport activities

Since R6G induces a sharp transition from the IF to the OF conformation over a narrow range of ATP-Mg^2+^ concentration, the effect on ATPase activity was investigated in this range. Indeed, when investigated at high ATP-Mg^2+^ concentration with R6G, Doxorubicin or Hœchst33342, no stimulation was observed [11, 32, 33]. However, at low ATP-Mg^2+^ concentrations, a clear stimulation is observed (Figure 4ABC), corresponding to the concentration range where the IF to OF transition occurs. Similar stimulation is observed in detergent, liposome or in membrane vesicles (Figure 4D). In all cases, where the ATP-Mg^2+^ concentration is above the K_d-app_ for ATP, i.e. ATP binding is not limiting anymore, no stimulation is observed confirming previous studies. We then investigated the transport efficiency for Hœchst33342 and Doxorubicin at varying ATP-Mg^2+^ concentrations (Figure 4E-F, Supp-Figure 23-24) as unfortunately R6G transport cannot be followed in membrane vesicles [13] but they nevertheless induce allosteric binding of ATP-Mg^2+^ (Supp-Figure 25). We quantified on the same membrane fraction the Hœchst33342 transport and ATPase activity of BmrA, to determine the amount of substrate being transported in relation to the ATP consumption. The experiments were carried out the same day for a precise quantification of both activities and conducted in quadruplicates from two separate batches of membrane preparation. Hœchst33342 transport per ATP hydrolyzed increases up to a maximum of 0.8 for 200-300 µM ATP-Mg^2+^, and slowly decreases afterwards. This biphasic curve implies that two phenomena are occurring. At low ATP-Mg^2+^ concentrations, a near strict coupling of Hœchst 33342 transport with ATP hydrolysis happens, while ATP gets hydrolyzed faster than its use for substrate transport at higher ATP-Mg^2+^ concentrations. For Doxorubicin, a similar curve is observed with a wider peak spreading over 300-1000 µM ATP-Mg^2+^, and reaching a maximum of 0.2 Doxorubicin transported per ATP hydrolyzed. This seems to also indicate that when ATP-Mg^2+^ concentrations are low resulting in slow binding to BmrA, the drug is transported more efficiently in regard to ATP consumption, while this effect disappears when the later concentrations rise.

**Figure 4:**
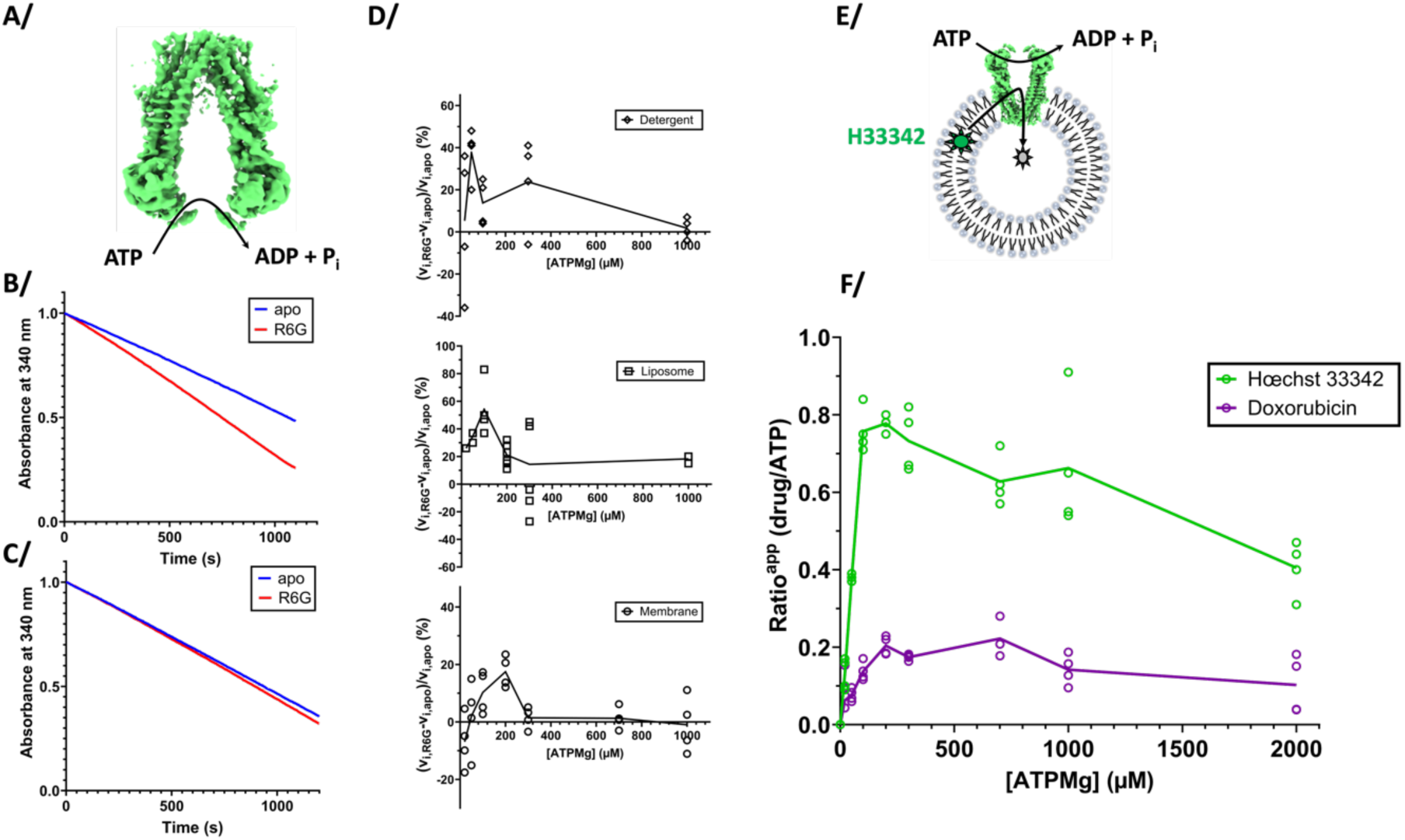
ATPase activities and transport measurements. **A/** Schematic ATPase activity of the transporter **B/** ATPase activity can be measured by following disappearance of ATP via a coupled enzymatic assay; decrease of absorbance at 340nm is directly correlated to ATP hydrolysis. Blue curve for BmrA apo, and red in complex with R6G. The slope is greater with R6G showing a faster ATP hydrolysis. This example is for the point 300 µM ATP for BmrA in detergent as shown in D/.**C/** same as B/ for 1 mM ATP showing no stimulation. **D/** ATPase activity measured in detergent, liposomes or in membranes. Each point represents a kinetic experiment as in B/ for a range of ATP concentration, acquired with or without R6G. The slope of each curve was measured at initial speeds, and the normalized ratio is presented on the graph to show the activation. **E/** Scheme of Hœchst 33342 transport in membranes occurring with ATP hydrolysis. **F/** Transport efficiency of Hoechst 33342 or Doxorubicin transported per ATP hydrolyzed along the range of ATP-Mg^2+^ sampled in C/. Individual measures are indicated on the graph (quadruplicates) and the line links the average of each concentration of ATP-Mg^2+^ sampled. Details of this experiment are shown in supp-Figures 23-24.

## 3. Discussion

Allostery between the substrate-binding site and the NBDs in ABC transporters has been very often described for many transporters and in various settings (a few examples here [7, 34–37]). It remains however difficult to study and to investigate at the molecular level being the result of multiple factors. Here we used the sharp effect produced by R6G on BmrA, to visualize the IF to OF transition, over a narrow range of ATP-Mg^2+^ concentration; the use of the ATPase inactive mutant E504A, which still binds ATP-Mg^2+^ but does not hydrolyze it, permits this structural enzymology approach as the mutant only undergoes the forward direction and is blocked in the OF conformation (Figure 1). The allostery manifests in several ways, visible at the NBD site. First, there is a structural rearrangement of the NBDs towards each other. This is possible with the intrinsic plasticity of BmrA in the IF conformation that allows to sample different conformations (Figure 2A-C), as also previously observed by NMR spectroscopy, HDX-MS or SANS studies [15–17]. Substrate binding to the drug-binding cavity thus optimizes the positioning of the NBDs. Second, the conformational space exploration of the NBDs is also influenced by R6G binding, as probed by the continuous heterogeneity analysis of cryoEM data (Figure 3). The substrate reduces the overall motions undergone by the NBDs, decreasing rotations and translations of the sites towards one another. This reduction in space exploration is also visualized by MD simulations where R6G also focuses the motions sampled by the NBDs (Supp-Figure 19). In both observations, the NBDs are still very mobile, the presence of the drug does not prevent movement but rather allosterically influences the directions of the space exploration. Since the ATP-binding sites (NBS) are shared between the two NBDs, ATP-binding implies that the NBDs come in close proximity. However, R6G prevents direct closure of the transporter using symmetrical closure as hypothesized for the apo WT transporter in absence of drug [21]; indeed, R6G binding in the drug-binding pockets prevents the TM helices to deform and to directly come close to the symmetric mates in the dimer. We hypothesize that this forces the TM helices to deform around the drug, which will have the consequence of forming a first NBS with ATP sandwiched between the Walker A motif of one NBD and the signature C motif of the opposite NBD (Figure 5). This site would correspond to the high affinity site and would help prime the other NBS for ATP binding, in accordance with the allosteric model. Of note, allostery at the NBS level during ATP hydrolysis has previously been observed by solid-state NMR for BmrA, illustrating that the two NBS can be asymmetrical of each other and reinforcing the plastic behavior of this transporter[11, 15].

**Figure 5:**
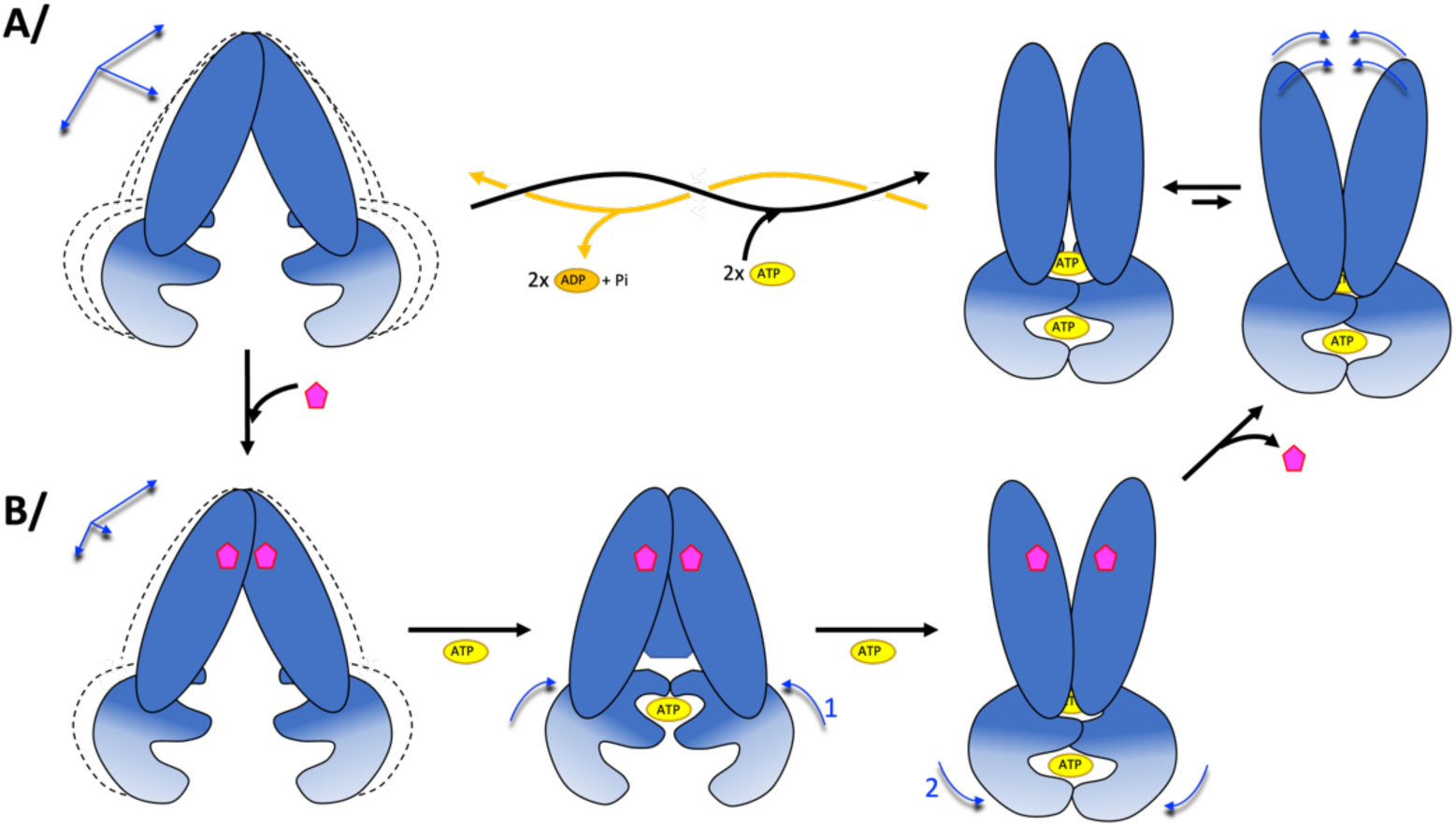
Model of transport mechanism. **A/** In absence of substrate, BmrA in the IF conformation is very plastic. The 2 NBS bind 2 ATP-Mg^2+^ (yellow cylinder) identically, leading to the OF conformation. Plastic deformation is transferred to the other side of BmrA for release of a potential substrate; return to the IF conformation is achieved via ATP hydrolysis and release of ADP-Mg^2+^ (orange cylinder). **B/** In presence of substrate, flexibility of the IF conformation is reduced and symmetrical closing is prevented by the presence of the drug. Binding of the first ATP-Mg^2+^ occurs by deformation around the drug, leading to the formation of the first NBS, which increases affinity for the other NBS and formation of the OF conformation. Using plasticity of the OF side of BmrA, substrate is released and the initial pathway of ATP hydrolysis is used to reset to the IF conformation.

Since the allosteric effect of R6G is to reduce the conformational space and to focus the movements explored by the NBDs, with a consequence of a sharp transition from IF to OF over a narrow range at ATP-Mg^2+^, we hypothesized that the drug would stimulate ATP hydrolysis in this narrow range of ATP-Mg^2+^ concentration. Indeed, such observation was made, in detergent, liposomes or membrane vesicles, with a peak of stimulation centered on 100-300 µM ATP-Mg^2+^, supporting the proposed model. However, the stimulation remains modest for BmrA compared to what can be observed for other Type-IV ABC transporters [38]. The maximum stimulation stays in the 30-40% activation, like what was also observed for reserpine, another substrate of BmrA[11]. This led us to investigate the stoichiometry of transport for BmrA of two substrates, Hœchst33342 and doxorubicin. They both induce allosteric ATP-Mg^2+^ binding like R6G (Supp-Figure 25) and their transport assays can be followed by fluorescence using inverted membrane vesicles[11, 13, 39]. For both substrates, a peak of substrate transported per ATP hydrolyzed is observed for the same range of 100-300 µM ATP-Mg^2+^ where stimulation occurs, as well as the transition IF to OF is maximal following ATP-Mg^2+^ binding. The ATP range for which the substrate stimulates ATP hydrolysis thus also corresponds to a more efficient drug transport (Figure 4DF). One can also note that Hœchst33342 and Doxorubicin are transported with a different ratio of ATP usage by BmrA. Hœchst33342 and R6G binding pockets are shared in the homolog human ABCB1 [40], thus, since there are 2 R6G bound on BmrA it is most probable that 2 Hœchst33342 or Doxorubicin also bind in BmrA substrate cavity (Supp-Figure 25). Since 2 ATP-Mg^2+^ are required to achieve the OF conformation and the ratio Hœchst 33342:ATP-hydrolyzed reaches values close to 1:1, it is likely that at the peak, 2 Hœchst33342 will be transported by cycle, resulting in an almost fully coupled activity. The smaller than 1 ratio also indicates that ATPase activity nevertheless occurs without any substrate bound to the drug-binding cavity or that the IF to OF transition occurs with only one Hœchst33342 bound. Also, at higher ATP-Mg^2+^ concentrations, ATP-Mg^2+^ binds faster and gets hydrolyzed faster than Hœchst is being transported, pointing rather to an uncoupling of the activities. Altogether, this would point to the diffusion of substrate inside the drug-binding pocket as a limiting factor for transport, as previously hypothesized by the kinetic selection model [41]. This hypothesis is reinforced with doxorubicin having a much lower ratio towards ATP. Since the LogP of these two drugs are different (3 for Hœchst33342, 0.8 for Doxorubicin as determined by ACD percepta), it implies a lower partition of Doxorubicin within the lipid bilayer or lower accessibility from the cytoplasm to reach the drug-binding site. Altogether, this clearly reveals a decoupling between the activities of drug transport and ATP hydrolysis for BmrA. We thus hypothesize that the diffusion rate of drugs into BmrA binding pockets is the limiting factor for transport, while ATP is being used constantly[38]; if a drug is bound while a cycle is being performed upon ATP-Mg^2+^ binding, it will be transported.

The allosteric effect of drugs on ATP-Mg^2+^ binding (Supp-Figure 25) is nevertheless meaningful at low ATP-Mg^2+^ concentrations as it results in a stimulation of drug transport efficiency (Figure 4C). Low ATP concentrations can be found in bacteria under early or stationary phase in rich or minimum media[42], starvation conditions[43], during sporulation or within persister cells[44]. In these conditions, ATP is scarce and precious, and cannot be wasted on a transporter working constantly; thus drug binding would have a positive influence on ATP-Mg^2+^ binding and will help the transporter to transport them more efficiently by more strictly coupling the two activities, thereby protecting the cells against xenobiotics. Under more favorable growing conditions, ATP concentrations rise and the need for an allosteric effect is diminished as the transporter functions faster[38].

Finally, the allostery described here is a derivative from the well-established Monod-Wyman-Changeux (MWC) model that apply to many systems [45]. Instead of stabilizing by ligand binding some clearly defined R and T states influenced by ligand-binding, we observe here that R6G indeed influences the conformation of the protein by reorienting the NBDs, reminiscent of the two states described in the MWC model, but more so modulates protein dynamics and conformational space exploration. This observation opens new avenues of investigations to understand structure/function relationship within proteins.

## 4. Materials & Methods

### 4.1 BmrA expression and purification

C43(DE3)ΔAcrB *E.coli* strain is transformed with the plasmid pET15b encoding for the BmrAE504A mutant fused to a 6-histidine tag in N-terminal. A transformed colony is incubated in 3 mL of LB media supplemented with 50 µg/mL of ampicillin for 7 h in 37 °C with shaking. 30 µL from this day culture are diluted in 1 L of LB media containing 50 µg/mL of ampicillin and incubated overnight at 22 °C with agitation. When OD^600^ reaches 0.6, BmrA overexpression is induced by adding 0.7 mM IPTG. Culture is then incubated for 5 h at 22 °C under shaking. Bacteria are collected by centrifugation at 5000 xg for 15 min, 4 °C. Pellet is then suspended in 20 mL of 50 mM Tris-HCl pH8.0, 5 mM MgCl_2_. Bacteria are lyzed by 3 passages with a disruptor Constant CellD system at 1.5 kbar, 4 °C. The bacterial suspension is centrifuged at 15,000 xg, 30 min, 4 °C. Supernatant is centrifuged for 1 h, 180,000 xg, 4°C to pellet membranes. The membrane fraction is suspended in 25 mL of 50 mM Tris-HCl pH8.0, 1 mM EDTA, anti-protease CLAPA 1X and centrifuged again with the same parameters. The membranes are finally suspended in 20 mM Tris-HCl pH8.0, 300 mM sucrose, 1 mM EDTA, frozen in liquid nitrogen and stored at -70 °C.

Membranes are solubilized at 3 mg/mL in 20 mM Tris-HCl pH 8.0, 100 mM NaCl, 15% glycerol (v/v), anti-protease CLAPA 1X, 4.5% (v/v) Triton X100 and incubated 1h20 at 4 °C under gentle agitation. The solution is centrifuged 40 min at 100,000 xg, 4 °C. The supernatant is loaded onto a Ni^2+^-NTA column pre-equilibrated with 20 mM Tris-HCl pH8.0, 100 mM NaCl, 15% (v/v) glycerol, anti-protease CLAPA 1X, 4.5% Triton X100, 20 mM imidazole. Resin is washed with 20 mM HEPES-NaOH pH 8.0, 100 mM NaCl, 20 mM imidazole, 1.3 mM DDM and 1 mM sodium cholate. Protein is eluted with the same buffer with 200 mM imidazole. Fractions of BmrA are pooled and diluted ten times in the same buffer as previously but without imidazole. The column is equilibrated with 20 mM HEPES-NaOH pH8.0, 100 mM NaCl, 20 mM imidazole, 1.3 mM DDM and 1 mM sodium cholate, and the protein solution is loaded again for a second affinity chromatography. After elution, fractions of BmrA are pooled (typically 10 mL) and concentrated on 50 kDa cutoff Amicon Ultra-15 device at 1,000 xg, 4 °C, until the volume reaches 500 µL. The solution is then injected on a Superdex 200 10/300 column (Cytiva) equilibrated with 20 mM HEPES-NaOH pH 7.5, 100 mM NaCl, 0.7 mM DDM and 0.7 mM sodium cholate.

### 4.2 CryoEM grids preparation and data collection

Purified BmrAE504A was concentrated to 4 mg/mL as described above. Defined concentrations of ATP-Mg^2+^ (25 µM, 70 µM, 100µM, 5 mM final concentrations) are added to a final concentration of BmrA of 25 µM to be applied on the cryoEM grid. For the grid containing Rhodamine6G (R6G), the ligand is also added to a final concentration of 100 µM followed by a 15 min. incubation on ice prior to ATP-Mg^2+^ addition. The mix was incubated 30 min. at room temperature to reach steady state before application on the cryoEM grid.

Ultra-Au 1.2/1.3 grids (Quantifoil) are glow discharged on air for 45 s at 30 mA (Emitech Glow Discharge). A volume of 3.5 µl of the mix is applied on freshly glow discharged grids at 20°C and 100% humidity using a Vitrobot Mark IV (Thermofischer). Excess liquid is blotted 4 s at blot force 0 and 0.5 s drain time before vitrification in liquid ethane. Data were collected on 2 Titan Krios at 300 kV equipped with a K3 direct electron detector (ESRF CM01 or Diamond eBIC), or a Talos Glacios equipped with a K2 detector (IBS, Grenoble). Data collection parameters are summarized in SuppTable 1.

### 4.3 cryoEM data processing

Data were processed using cryosparc v2, v3 and v4 (Structura Bio) over the duration of the project, with the first data collected in Sept. 2019. Movies were submitted to patch motion correction and patch CTF estimation jobs, and particle picking was performed using blob picker (100-200 Å diameter) on a small subset of movies (usually 200) to create initial 2D classes to be used for template picking. Automatic particle picking was then performed on all the movies using the template from previous 2D classification. Mild 2D classification was operated to remove obvious bad particles (ice contamination or detergent micelles) but care was taken to not throw out “broken particles” at this stage. 3D maps are created with *ab-initio* reconstruction job asking for 7 models to observe the whole sample heterogeneity. The number of particles was noted for IF and OF classes to calculate their ratio (Figure 1). Each map was then individually further refined using several rounds of hetero-refinement and non-uniform refinement until the highest resolution was reached. Many routes were explored to reach high resolution reconstructions, leading to the final one displayed. Each job was computed with or without C2 symmetry to evaluate the impact on the maps; this is especially important for the datasets containing R6G to ascertain its density. In some cases, particle expansion followed by local refinement was performed to enhance the quality of the maps at the R6G binding site without imposing symmetry. The general data processing pipe was similar but some specificities were used for some datasets, to reach the best results possible. Summaries of data processing are shown in supp-Figures 1-8. Notably, for the data set E504A^apo-100µMATP^ an additional local CTF refinement was performed and a mask without detergent belt was applied. For data set E504A^apo-25µMATP^ a mask without detergent belt and one NBD was used. For datasets showing discrete heterogeneity but where one conformation couldn’t be resolved to high resolution, the reconstructions clearly show in which category (IF or OF) the transporter belongs (E504A^apo-25µMATP^, E504A^R6G-25µMATP^, E504A^R6G-70µMATP^). These reconstructions were used to count the number of particles and thus derive percentages of populations, but no atomistic model was built.

### 4.4 Model building

For the outward facing conformations, models were built using the previous OF structures of BmrA in absence or presence of R6G (PDB: 6R72 and 6R81 [13]). Several cycles of manual building in Coot and real-space refinement using phenix.refine were performed. Final models were checked for rotamer and Ramachandran outliers.

For IF conformations, the first reconstruction was achieved before the opening of the AlphaFold database. The model was built using the OF conformation, with manual deformation of trans-membrane helices to match the density, using coot and ISOLDE, and several rounds of refinement in Phenix until a model was correctly built. Shortly after, the AlphaFold database was released and the BmrA model, which was generated in IF conformation, was also used for comparison and model building. It turns out that the conformation created by AlphaFold is more closed compared to the IF reconstruction experimentally obtained, thus distortions of that model was also needed. It was also used for comparison with our model building to ascertain the registry, which was identical in both models (ours and AlphaFold), and comforted us in residue assignment. For the other models in IF conformation, local modifications were performed by cycles of manual building and refinement as described before. As discussed below, the NBDs undergoes significant movement in the IF conformation, and thus lacks clear electron density for the outmost regions. The core of the NBDs however is clearly defined, with key residues around W413 for example well defined and unambiguous side-chain assignment and positioning. For the outmost parts, the NBDs were placed following the clear model in the OF conformation. Final models and maps were deposited in the PDB and EMDB under the accession code listed in Supp-Table1.

### 4.5 Variability analysis and variability refinement

For E504A^apo^ and E504A^R6G^, 3D Variability Analysis (3DVA, Cryosparc) was performed on the particles from the last non-uniform refinement. Resolution was filtered at 6 Å for 3DVA calculation and display for E504A^R6G^, and at 4.5 Å for E504A^apo^. Masks from Non-uniform refinement were initially dilated by 20 Å. The 3D variability display job was performed in simple or intermediate modes with similar results. 5 components were asked as output but the main movements are observed in the 3 first components, which were subjected to variability refinement. Each 20 maps originating from 3DVA were input to variability refinement (phenix.varref [18]) along with the final refined model. 50 models were generated for each map, with restraints adapted to the resolution of 3DVA calculations, and only the best model was kept for each map. The resulting file is a model file containing 20 models with atomic coordinates fitted to each map, allowing for a visualization of the model in ChimeraX for detailed analysis. Movies were generated with ChimeraX.

### 4.6 Molecular dynamics simulations

For each condition E504A^apo^ and E504A^R6G^, ATP-Mg^2+^ was added on the NBD in its binding-site by superposing the Outward-Facing conformation of BmrA (PDB: 6r82). Proteins were then inserted into a bilayer using GROMACS (??) and equilibrated … MD simulation were carried out using Amber36 (??) force field for 700ns and performed in triplicate.

For each condition E504A^apo^ and E504A^R6G^, ATP-Mg^2+^ was added on the NBD in its binding-site by superposing the Outward-Facing conformation of BmrA (PDB: 6r82). MD simulations were performed for these models using the Amber 14:EHT force field implemented in the MOE software [46]. A lipid bilayer surrounding the transporter composed of 1,2-dioleoyl-sn-glycero-3-phosphoethanolamine and 1,2-dioleoyl-sn-glycero-3-phospho-rac-1-glycerol with a respective ratio of 3:1 was generated with the MOE lipid generator, using the PACKMOL-Memgen approach [47–49] TIP3P water molecules [50, 51] were added, as well as Na^+^ and Cl^−^ ions in order to simulate a 0.1M ionic force, and the pH was set at 7.4. Standard Amber 14:SB parameters were used for MD in amber version 22 suite software [52]. A quick minimization was performed prior to the start of the MD simulations. The geometry of all the molecules, bonds, and charges was then checked using the structure preparation tool in MOE, in order to start the molecular simulations. After a preliminary equilibration phase, 700 ns NTP molecular simulations were performed in triplicate. A frame was saved every 200ns, leading to a total of 3500 frames for each simulation. During MD simulation temperature (300K) a pressure (1 atm) were kept constant (NTP).

### 4.7 Ligand binding

R6G, Doxorubicin, Hœchst33342 and Vincristine binding were carried out by probing the intrinsic fluorescence change as a function of ligand concentration. Fluorescence was recorded on a SAFAS Xenius spectrophotofluorimeter set up at a constant photo multiplicator voltage of 570 V. Tryptophan residues or N-acetyl tryptophan amide (NATA) used as negative control were excited at 290 nm, and their fluorescence emission spectra were recorded between 310 and 380 nm, with a 5-nm bandwidth for excitation and emission. Experiments were done in a quartz cuvette in a final volume of 200 µL, in which increasing amounts of ligand were added. Resulting emission curves were integrated and deduced from the same experiments carried out with NATA, used at the same concentration than that of BmrA tryptophan residues. Data were plotted as a function of ligand concentration.

### 4.8 Reconstitution into liposomes

300 µL of *E. coli* total lipid extract (Avanti Polar lipids) in chloroform at 25 mg/mL were placed into a 50-mL glass balloon and evaporated using a gentle nitrogen stream while turning the balloon to create a monolayer, under a hood. The monolayer was resuspended and solubilized by addition of 300 µL of lipid buffer (20 mM HEPES-NaOH pH 7.5, 100 mM NaCl) supplemented by 75 µL DDM at 10 % (w/v), and vortexing at high speed for 5 minutes. The balloon was then placed on a gentle rocker at room temperature for 1h. BmrA purified in the DDM/Cholate mixture was added to 100 µL of solubilized lipids to ensure a molar ratio 1 BmrA dimer per 3000 lipids (considering an average MW_lipids_ = 750 g/mol, *e.g.* 40 µL of BmrA at 2.8 mg/mL). The solution was completed to 500 µl with lipid buffer and incubated 45 min. at room temperature under gentle agitation. Detergents were then removed by 3 additions of 40 mg pre-activated biobeads, each followed by an incubation of 1h at room temperature under gentle agitation between each biobead addition. The final proteoliposomes solution was then extruded by passing 11 times through a 400-nm membrane filter (Avanti Polar Lipids). These final proteoliposomes were kept at 4°C and used immediately. BmrA retains its activity during 1 week in these conditions.

### 4.9 ATP hydrolysis assay

The ATPase activity of BmrA was measured as previously described[13]. The protein in solution in 20 mM HEPES-NaOH pH 7.5, 100 mM NaCl, 0.7 mM DDM and 0.7 mM Na cholate was diluted in the ATPase activity assay buffer containing a mixture of 0.7 mM DDM and 0.7 mM Na cholate, and the ATPase activity recorded. ATP hydrolysis was measured using an enzymatic coupled assay, where ADP is regenerated in ATP by the Pyruvate kinase at the expense of PEP and generating pyruvate. Pyruvate is reduced into lactate by the Lactate dehydrogenase by oxidizing NADH into NAD+. The disappearance of NADH was followed by absorbance at 340nm, indicative of ATP hydrolysis. Initial and maximum velocity were recorded. To represent the increase in velocity, the difference between velocities in presence and absence of R6G were normalized to the velocity in absence of R6G. ATPase activities were conducted at fixed ATP-Mg^2+^ concentration and varying R6G concentrations, or at fixed R6G concentration and varying ATP-Mg^2+^ concentrations. All ATP hydrolysis assays were realized with 4 µg wild-type and E504A inactive mutant in different amphipathic environments (detergent, liposome, inverted membrane vesicles). R6G was pre-incubated with the protein at 50 µM for 10 min at 25°C.

### 4.10 Transport experiments

Transport assays followed the fluorescence of Hœchst 33342 (Sigma-Aldrich) or doxorubicin (Sigma-Aldrich) during their transport by BmrA across the membrane with different ATP-Mg^2+^ (Sigma-Aldrich) concentrations. The set-up experiment consisted of 50 µg *E. coli* inverted membrane vesicles containing overexpressed BmrA mixed with 1 micromolar of fluorescent molecules (Hœchst 33342 or doxorubicin, stock solution resuspended in water). Transport was initiated by adding a certain concentration of ATP-Mg^2+^ and monitored at 25 °C in 1 mL quartz cuvettes, recording the fluorescence on Xenius fluorimeter (SAFAS) at 468 nm with a bandwidth of 10 nm upon excitation at 355 nm with a bandwidth of 10 nm for Hoechst 33342 and at 593 nm with the bandwith of 10 nm upon excitation at 468 nm with the bandwith of 10 nm. Different ATP-Mg^2+^ concentrations were tested, of 20, 50, 100, 200, 300, 700, 1000 and 2000 µM. The transport buffer was made of 50 mM HEPES-NaOH pH = 7.5 (Sigma-Aldrich), 8.5 mM NaCl (Sigma-Aldrich), 5 mM MgCl_2_ (Sigma-Aldrich), supplemented by 5 mM NaN_3_ (Sigma-Aldrich) and 2 mM Na_2_S (Sigma-Aldrich) which are inhibitors of electron transport chain. Transport assays with E504A inactive mutant were also performed to visualize some potential non-specific transport. All experiments were done in duplicate of duplicates. Standard curves were performed to estimate the amount of drug transported. They were performed on the same inverted membrane vesicles, or in pure liposomes with a fixed lipid concentration (The liposomes concentration was set to reach the same maximum fluorescence for the ligand as in membrane vesicles, which resulted in 10 mg/ml lipids). The curves were performed from zero to the maximum amount of substrate added, in small increments.

## Acknowledgements

We acknowledge the European Synchrotron Radiation Facility for provision of beam time on CM01 and we would like to thank all the staff for assistance. We thank FRISBI (PID-169 and - 202) and INSTRUCT (PID-24618) for funding access to microscopes. This work used the platforms of the Grenoble Instruct-ERIC center (ISBG; UAR3518 CNRS-CEA-UGA-EMBL) within the Grenoble Partnership for Structural Biology (PSB), supported by FRISBI (ANR-10-INSB-05-02 & project ID 160 to P.B.) and GRAL, financed within the University Grenoble Alpes graduate school (Écoles Universitaires de Recherche) CBH-EUR-GS (ANR-17-EURE-0003). The electron microscope facility is supported by the Auvergne-Rhône-Alpes Region, the Fondation pour la Recherche Médicale (FRM), the fonds FEDER and the GIS-Infrastructures en Biologie Santé et Agronomie (IBiSA). We thank Daouda Traoré for personal time on the ESRF CM01 Titan Krios. This project was funded by ANR-19-CE11-0023-01 for VC, PF, AG, SM, CO and JMJ, and ANR-23-CE11-0031-01 for VC, PF, SM, LM, LZ and GS. The authors wish to thank Elise Kaplan for sharing models of BmrA WT and C582, and Alexis Michon with his constant IT help during the course of this article.

## Author contribution

VC initiated the study. AG and LM expressed all the proteins, purified them and prepared them for cryoEM data acquisition and enzymologic characterization. AG processed all the data for the apo form and R6G-bound proteins, constructed all the models and performed 3DVA calculations and movies. AG performed the ATP binding assay for E504^apo^ and in presence of R6G. LM performed all the other biochemical and enzymologic analysis. SM prepared protein and membranes. EZ and GS prepared all the grids and observed them on the IBS GLACIOS with overnight data collection for 2D classes, and performed some data collection on the CM01 KRIOS. CO and JMJ performed early analysis on ATPase stimulation by drugs at low ATP concentrations. EK, CO and JMJ shared the WT and C582 structures. EB and RT performed MD simulations and logP analysis. JM performed analysis of the varref bundles. VC performed the variability refinement analysis. PF and VC supervised the whole project. All authors participated in writing the manuscript.

## Supp-Data

**Supp-Figure 1:**
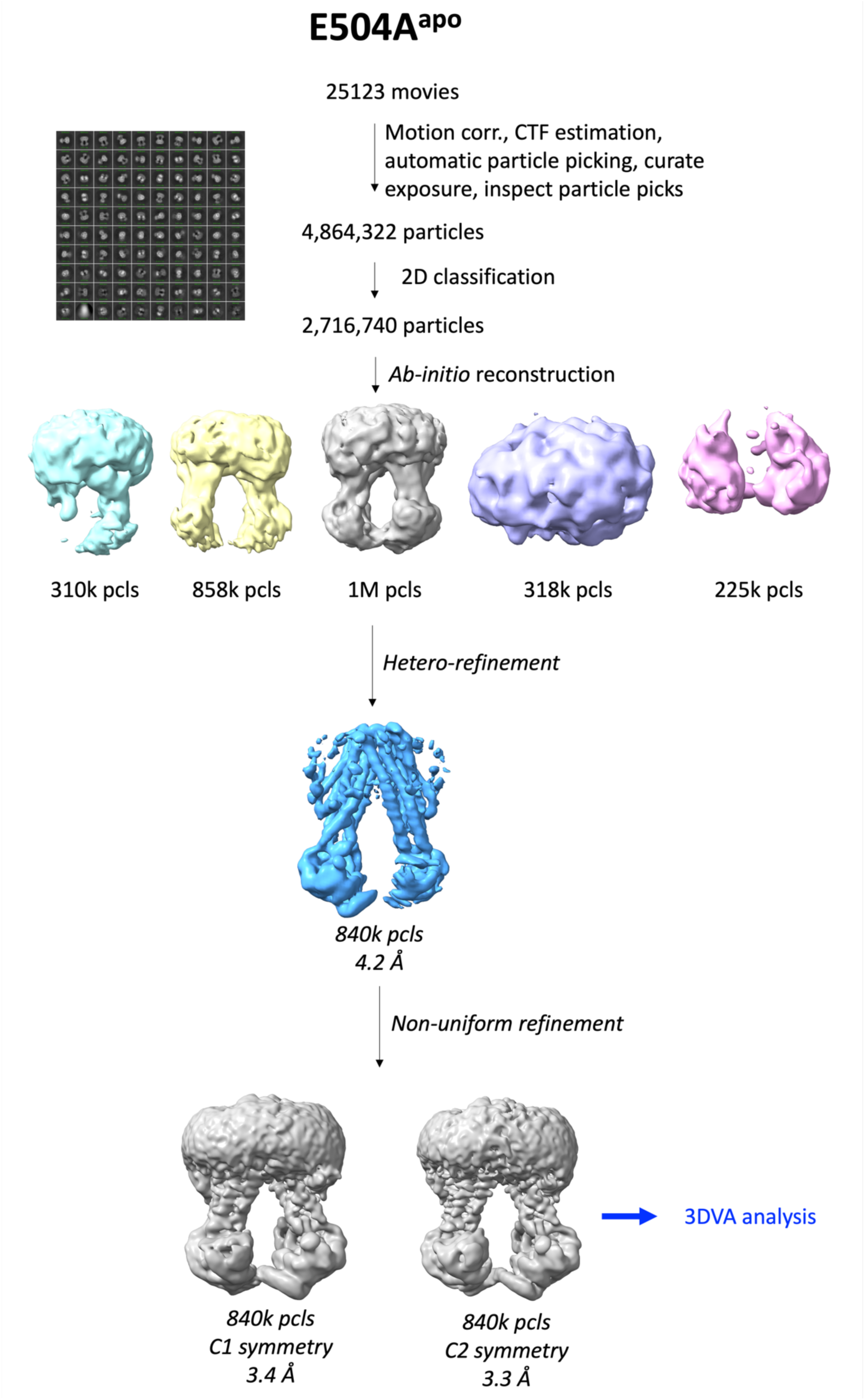
Data processing scheme for BmrA E504A^apo^. Particles (pcls) are listed for each step and class, and resolution at FSC^=0.143^ are listed for the latest stages of refinement. Many routes were explored to reach high resolution reconstructions, only the final one is displayed.

**Supp-Figure 2:**
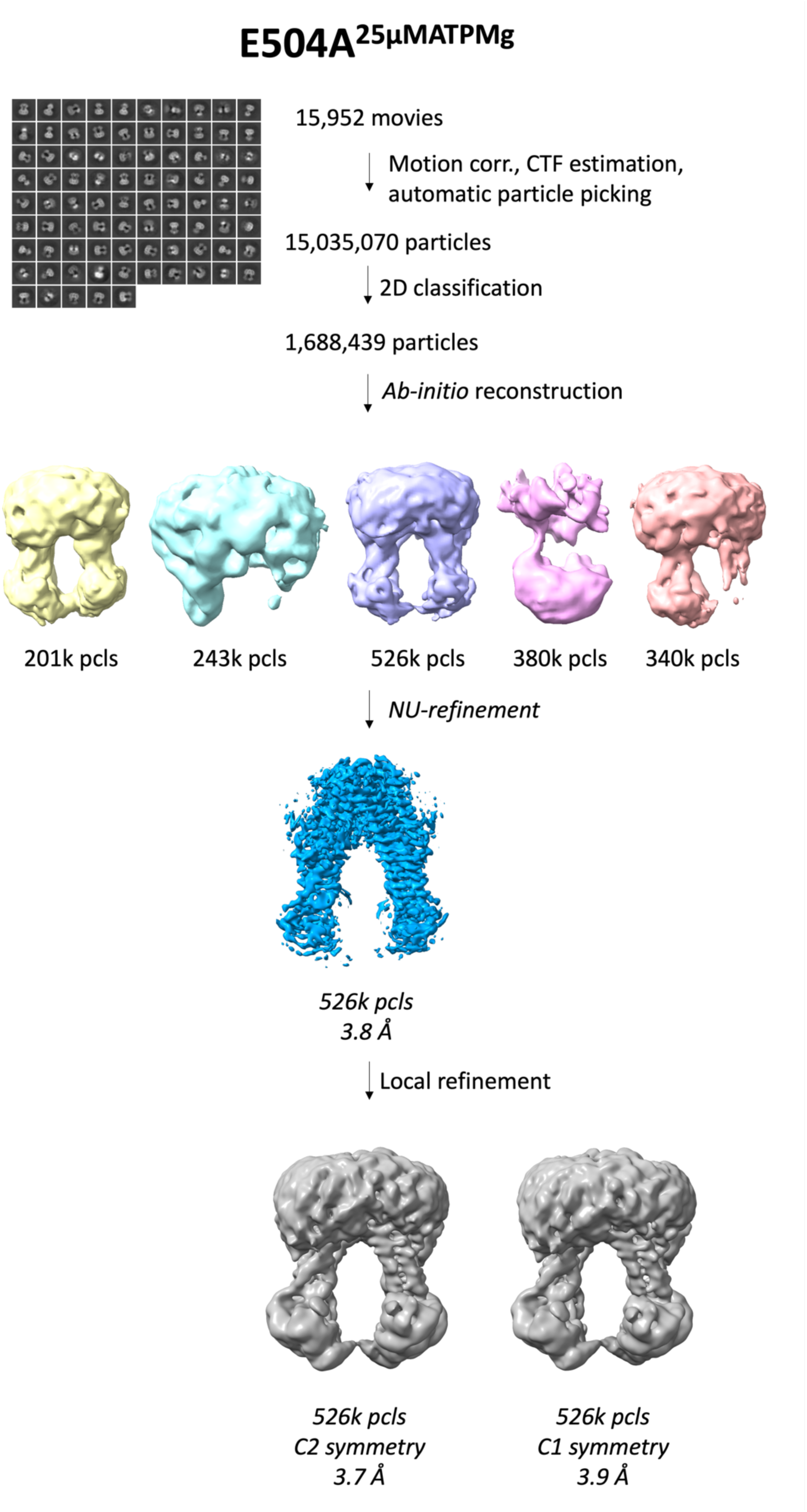
Data processing scheme for BmrA E504A_25μMATPMg_. Particles (pcls) are listed for each step and class, and resolution at FSC_=0.143_ are listed for the latest stages of refinement. Many routes were explored to reach high resolution reconstructions, only the final one is displayed.

**Supp-Figure 3:**
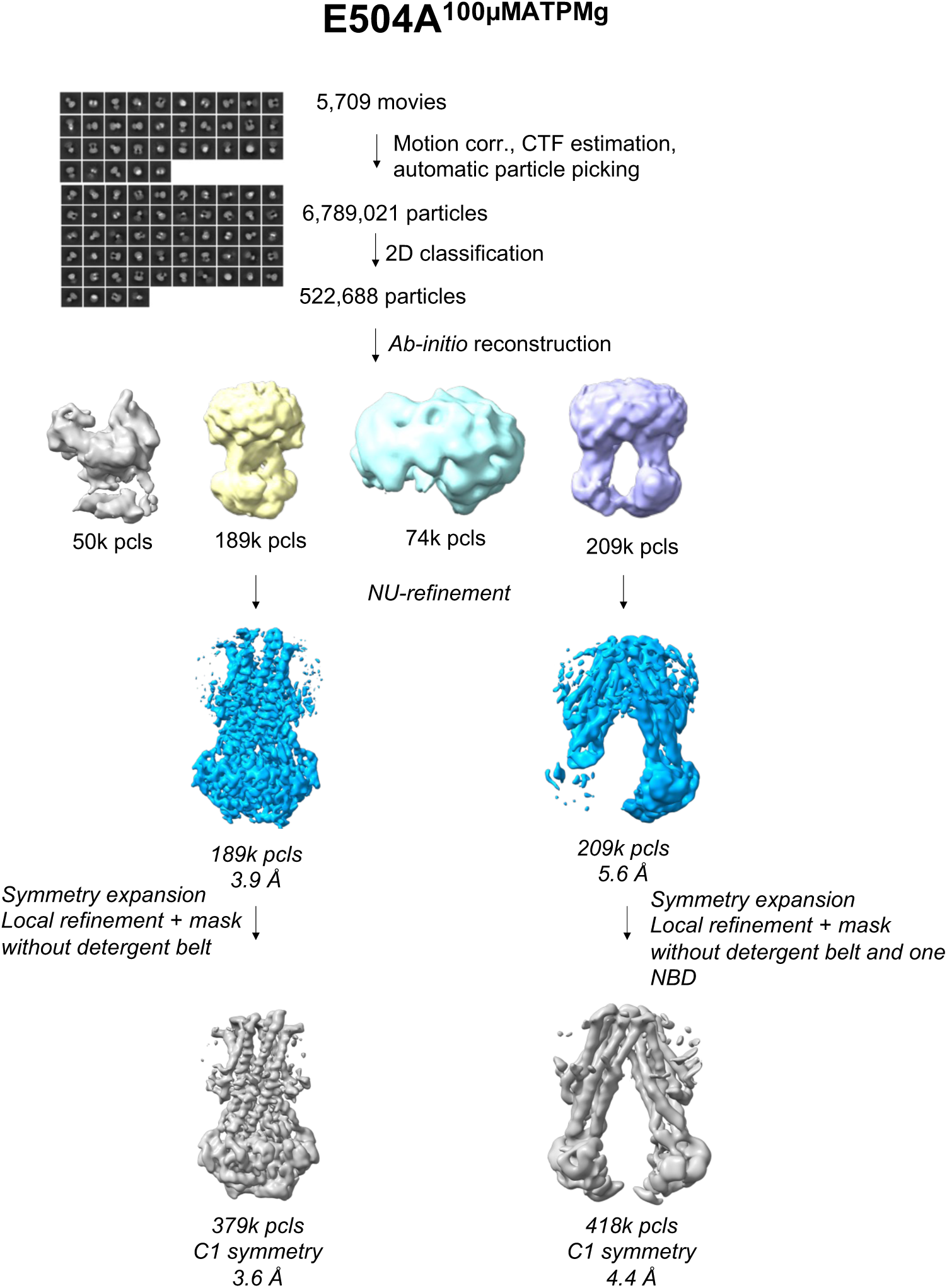
Data processing scheme for BmrA E504A^100µMATPMg^. Particles (pcls) are listed for each step and class, and resolution at FSC^=0.143^ are listed for the latest stages of refinement. Many routes were explored to reach high resolution reconstructions, only the final one is displayed.

**Supp-Figure 4:**
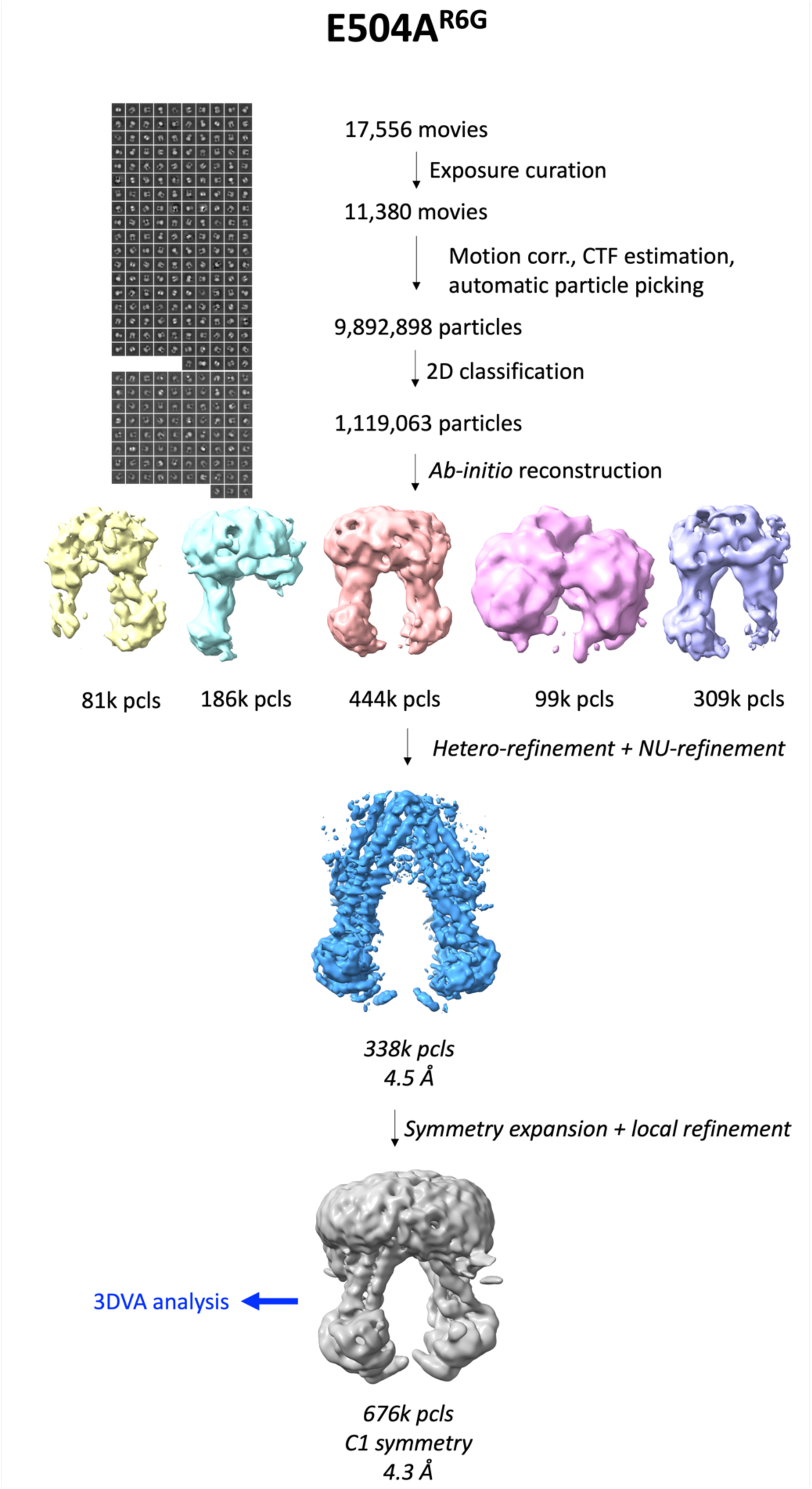
Data processing scheme for BmrA E504A^R6G^. Particles (pcls) are listed for each step and class, and resolution at FSC^=0.143^ are listed for the latest stages of refinement. Many routes were explored to reach high resolution reconstructions, only the final one is displayed.

**Supp-Figure 5:**
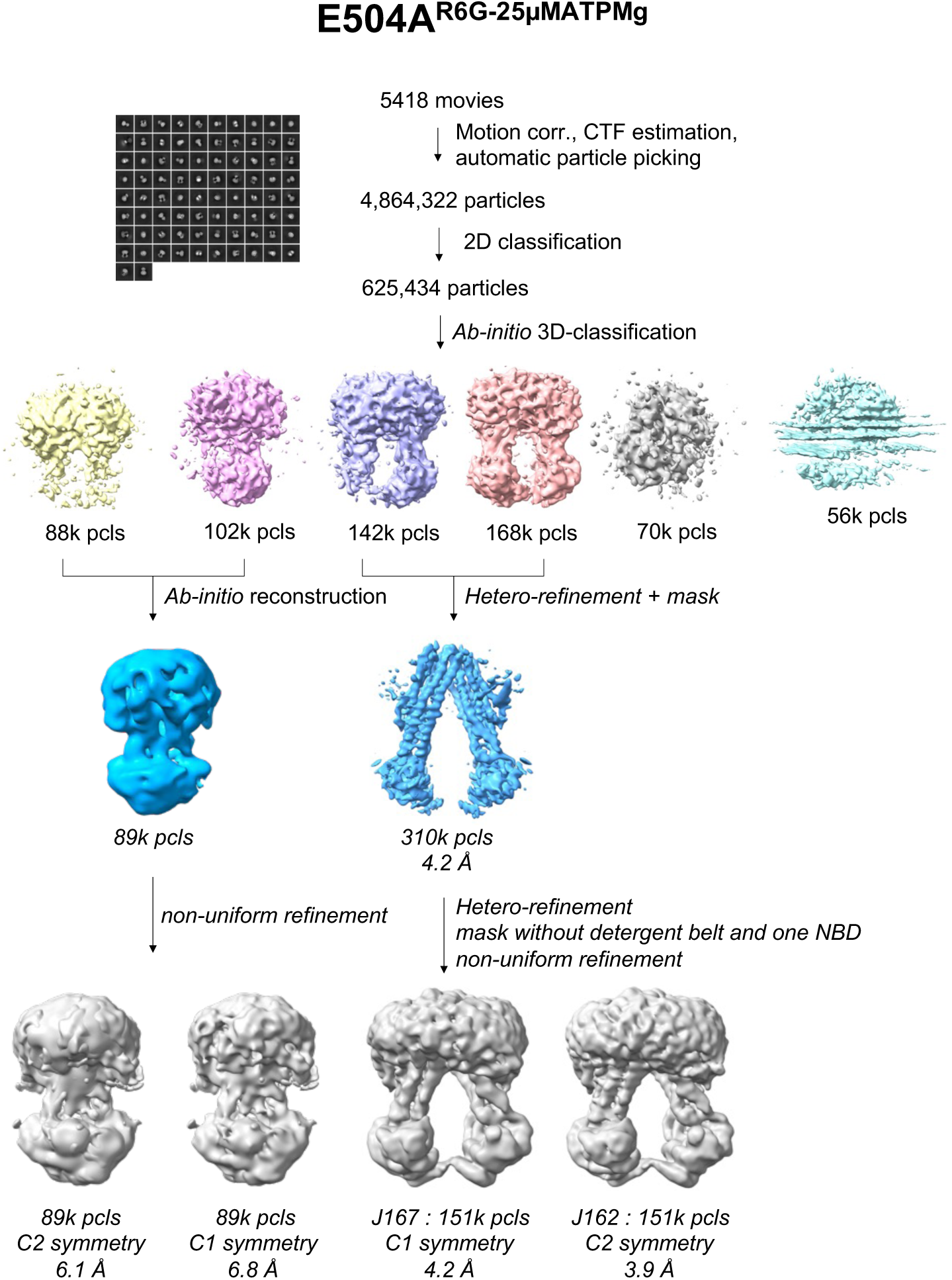
Data processing scheme for BmrA E504A^R6G-25μMATPMg^. Particles (pcls) are listed for each step and class, and resolution at FSC^=0.143^ are listed for the latest stages of refinement. Many routes were explored to reach high resolution reconstructions, only the final one is displayed.

**Supp-Figure 6:**
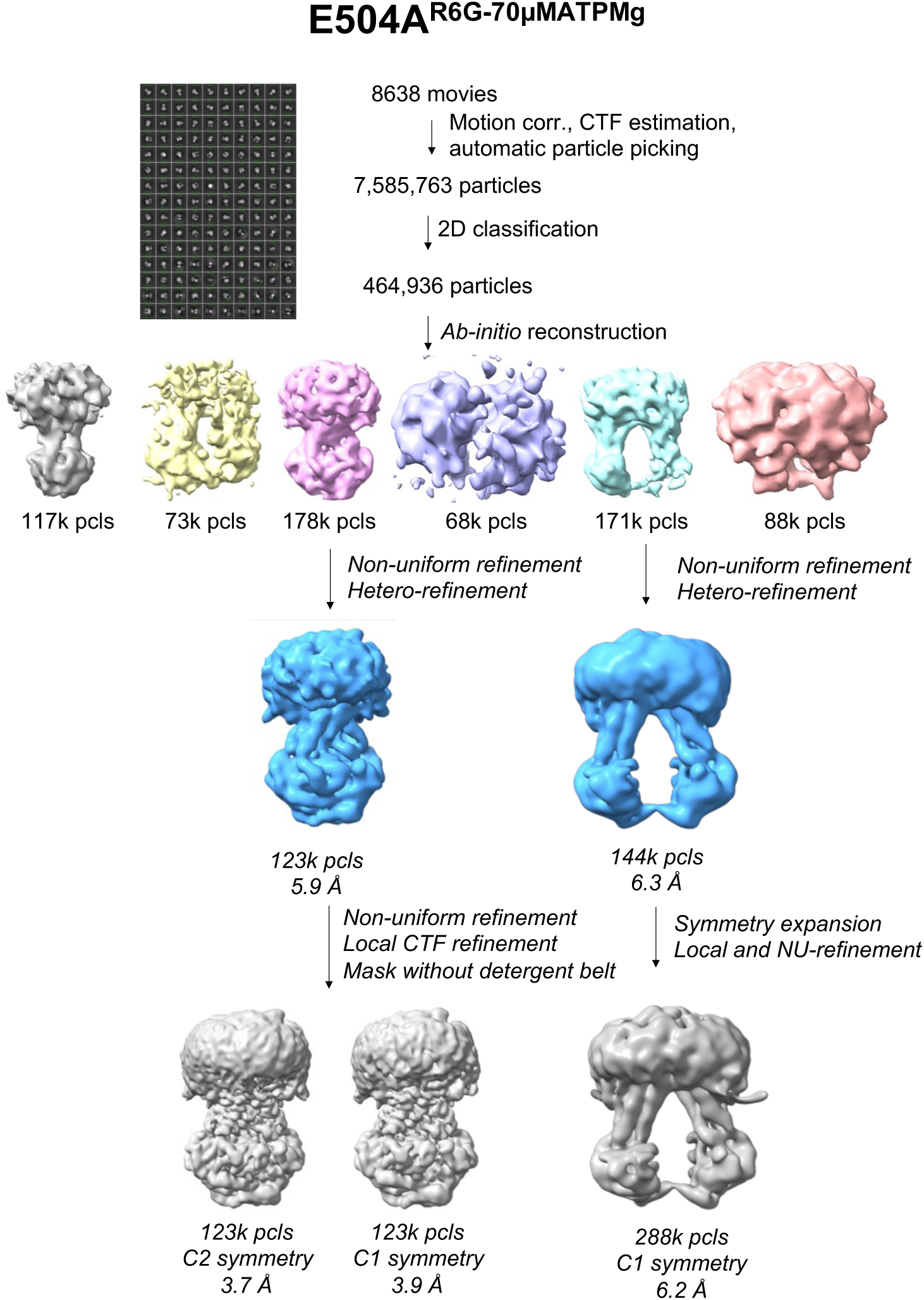
Data processing scheme for BmrA E504A^R6G-70µMATPMg^. Particles (pcls) are listed for each step and class, and resolution at FSC^=0.143^ are listed for the latest stages of refinement. Many routes were explored to reach high resolution reconstructions, only the final one is displayed.

**Supp-Figure 7:**
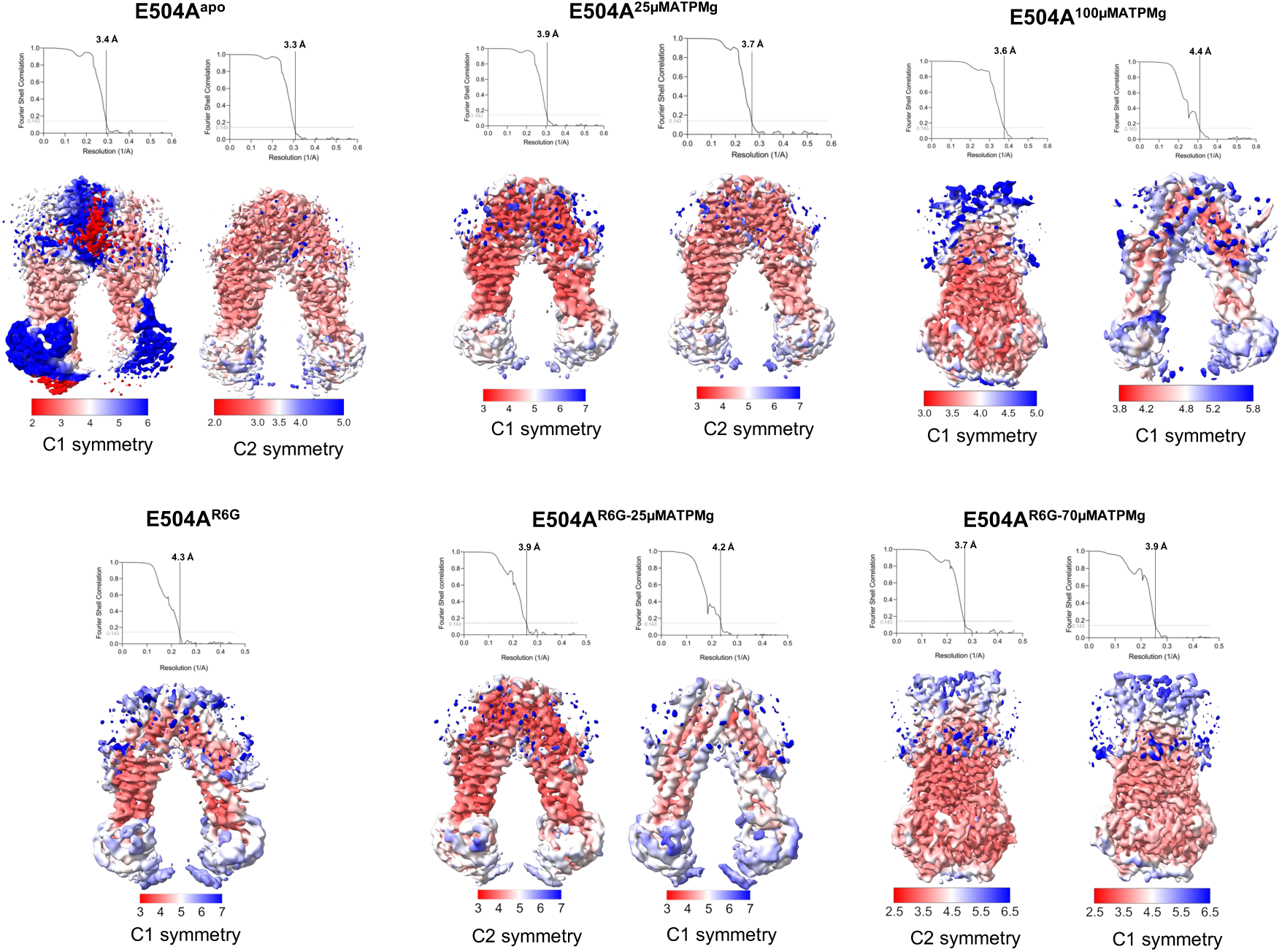
Local resolutions estimations for each dataset. Without R6G on top, with R6G at the bottom. Resolution scales vary for each dataset to better render the resolution range of each reconstruction.

**Supp-Figure 8:**
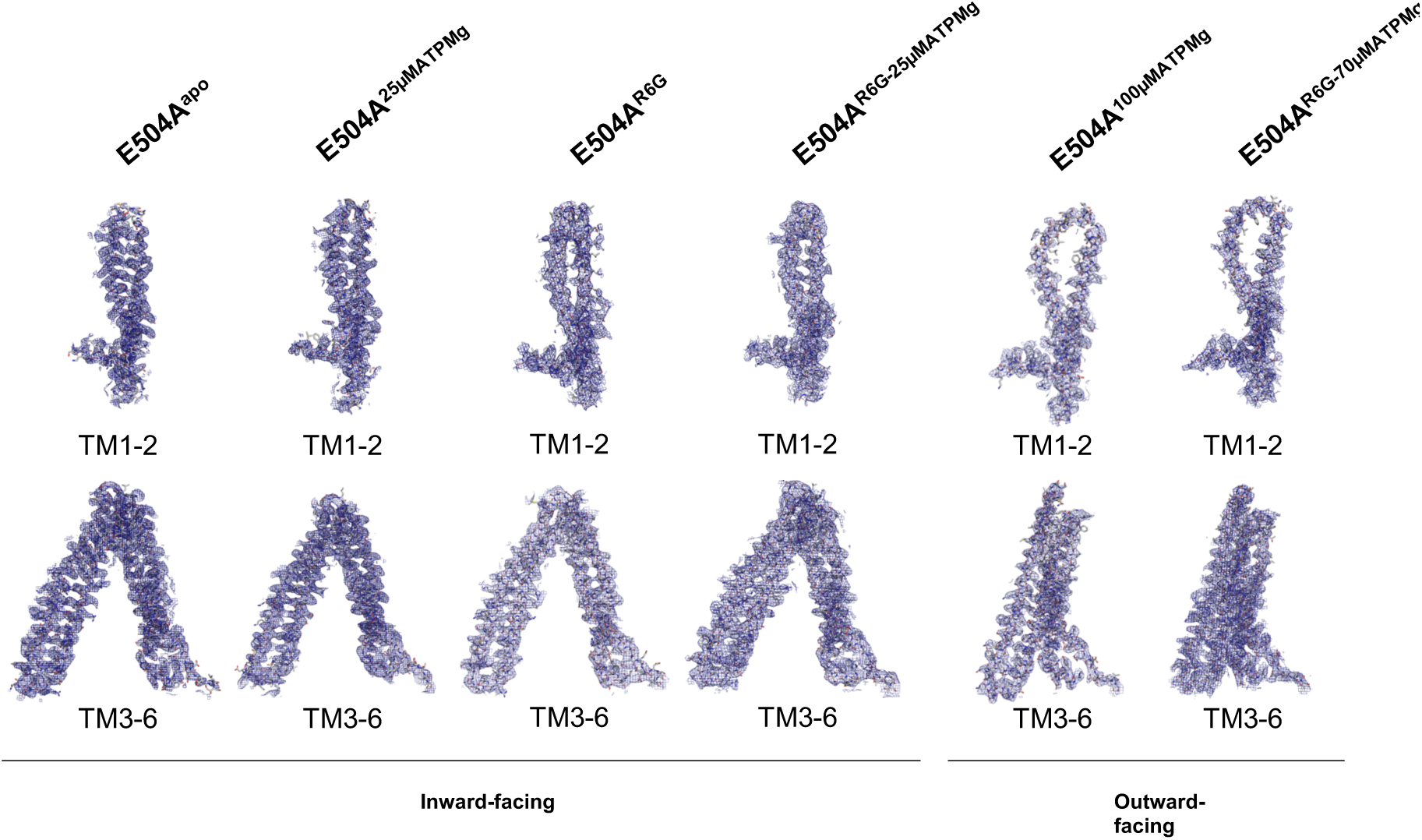
Coulomb potential maps for the trans-membrane regions of each dataset, exemplifying the backbone and side chains can be placed unambiguously and that the conformational changes induced by substrate binding are indeed reflected by changes in reconstructions.

**Supp-Figure 9:**
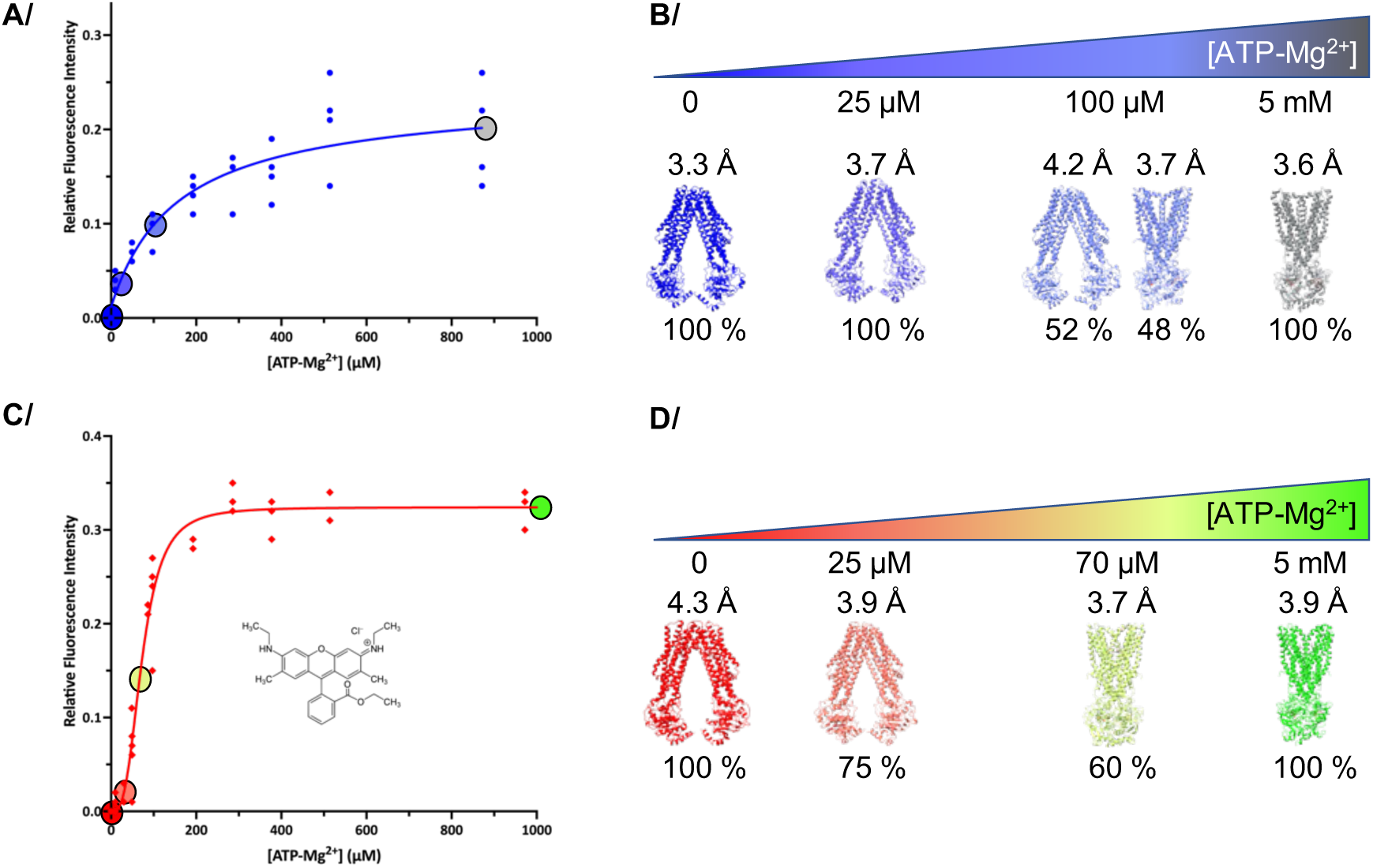
Models generated for each reconstruction. **A/**, **B/**, **C/** and **D/** as in Figure 1, with the same color code. Models are represented instead of reconstructions in **B/** and **D/**, when resolution was sufficient to generate a model.

**Supp-Figure 10:**
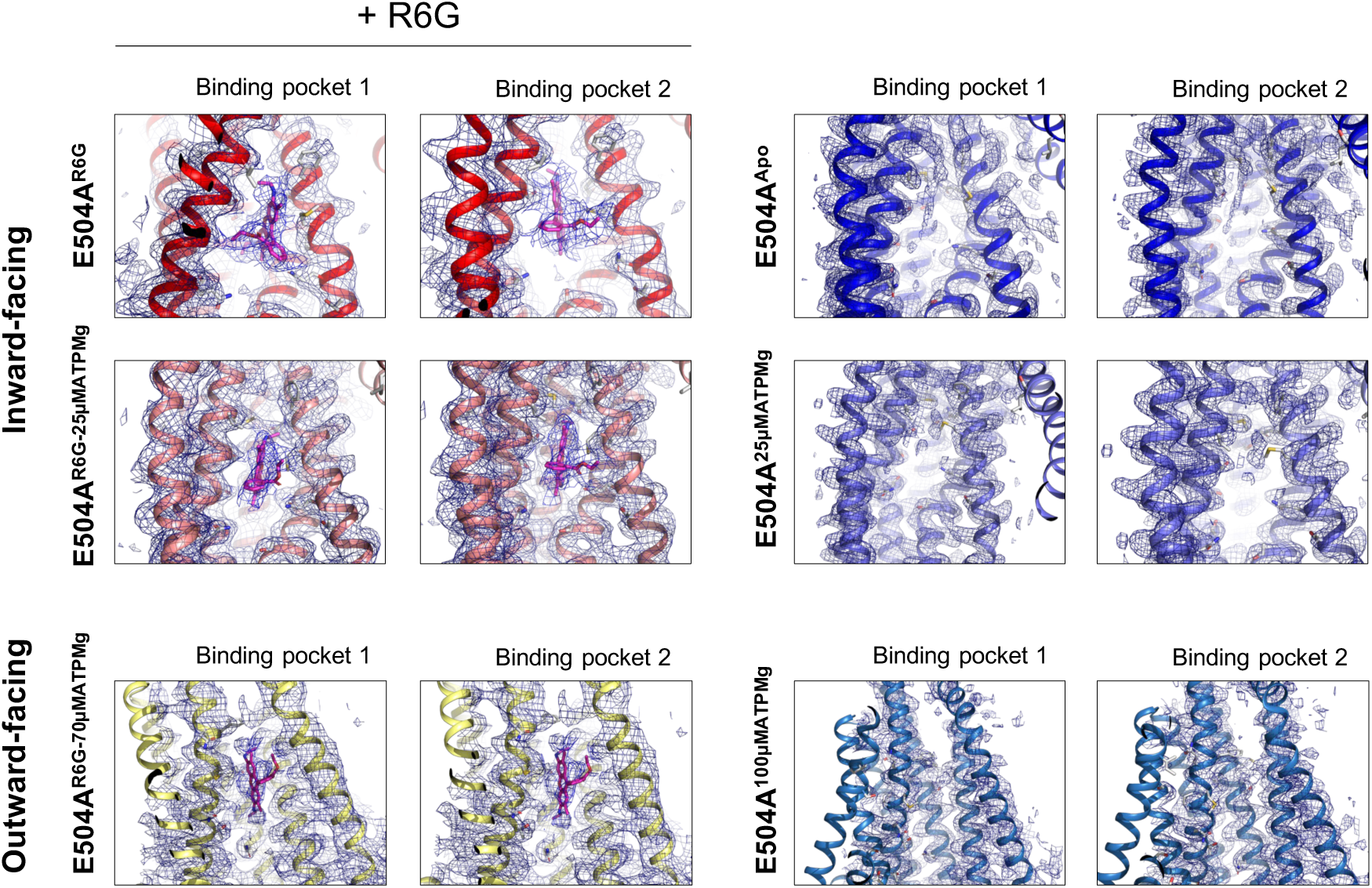
Coulomb potential maps for R6G in the binding pocket. Left side displays all datasets with R6G present in the solution, right side without R6G. Maps are shown in blue mesh, and the models are displayed in cartoon with the same coloring scheme as in Figure 1. For each dataset, the two halves of BmrA are displayed (binding pockets 1 and 2). R6G is shown as magenta sticks, and its density is indeed present each time R6G is added to the protein.

**Supp-Figure 11:**
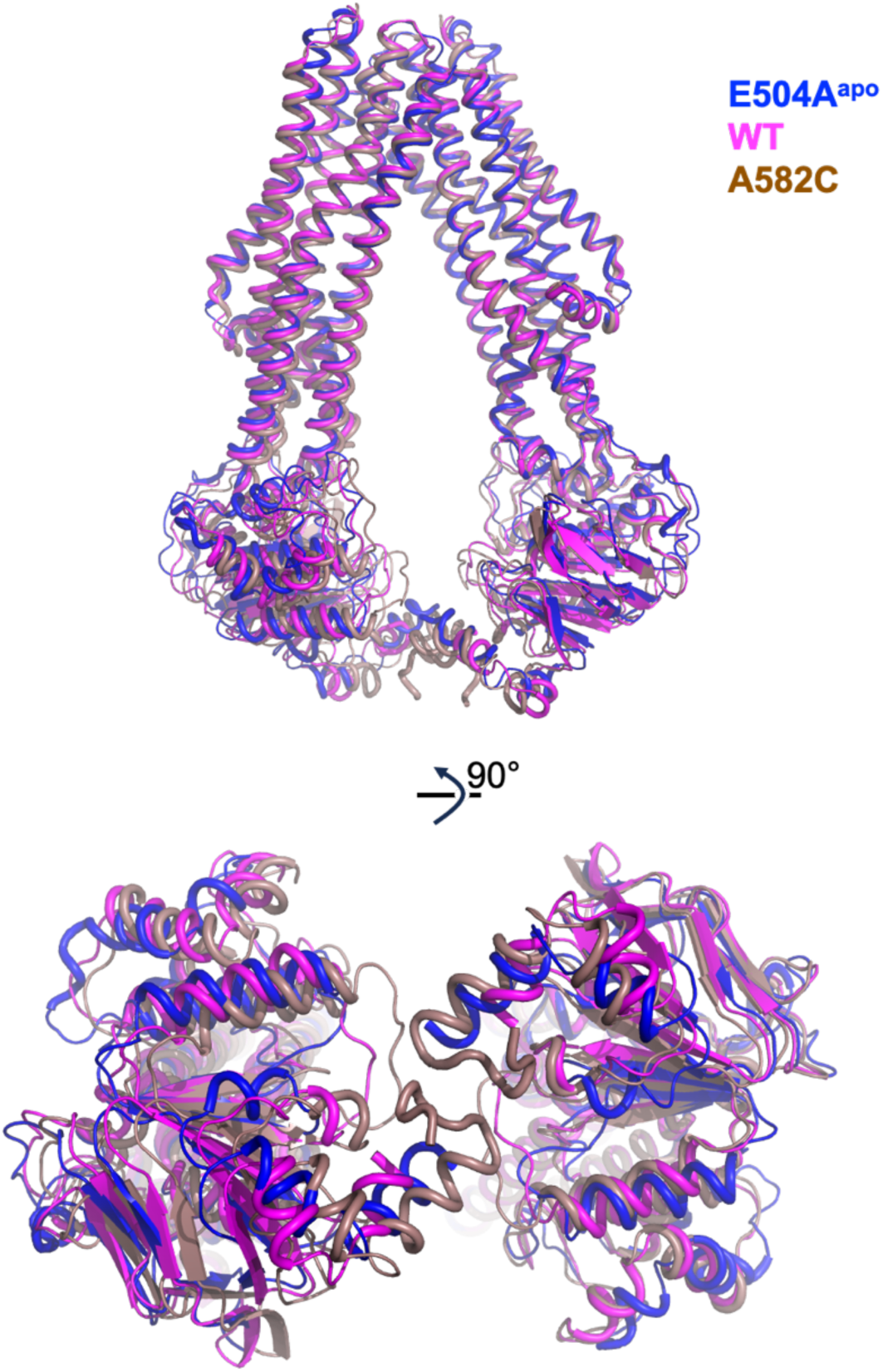
Overlay of BmrA WT, A582C and E504A in the apo forms. E504A^apo^ is colored in blue according to the color scheme of Figure 1. BmrA WT is colored magenta, and the A582C mutant is colored brown. All proteins are represented in cartoon from the side (top) and from the NBDs (bottom). Overlay performed on residues 1-161 of chain A for each structure with a rmsd of 0.52Å over 902 atoms and 0.45 Å over 883 atoms, respectively.

**Supp-Figure 12:**
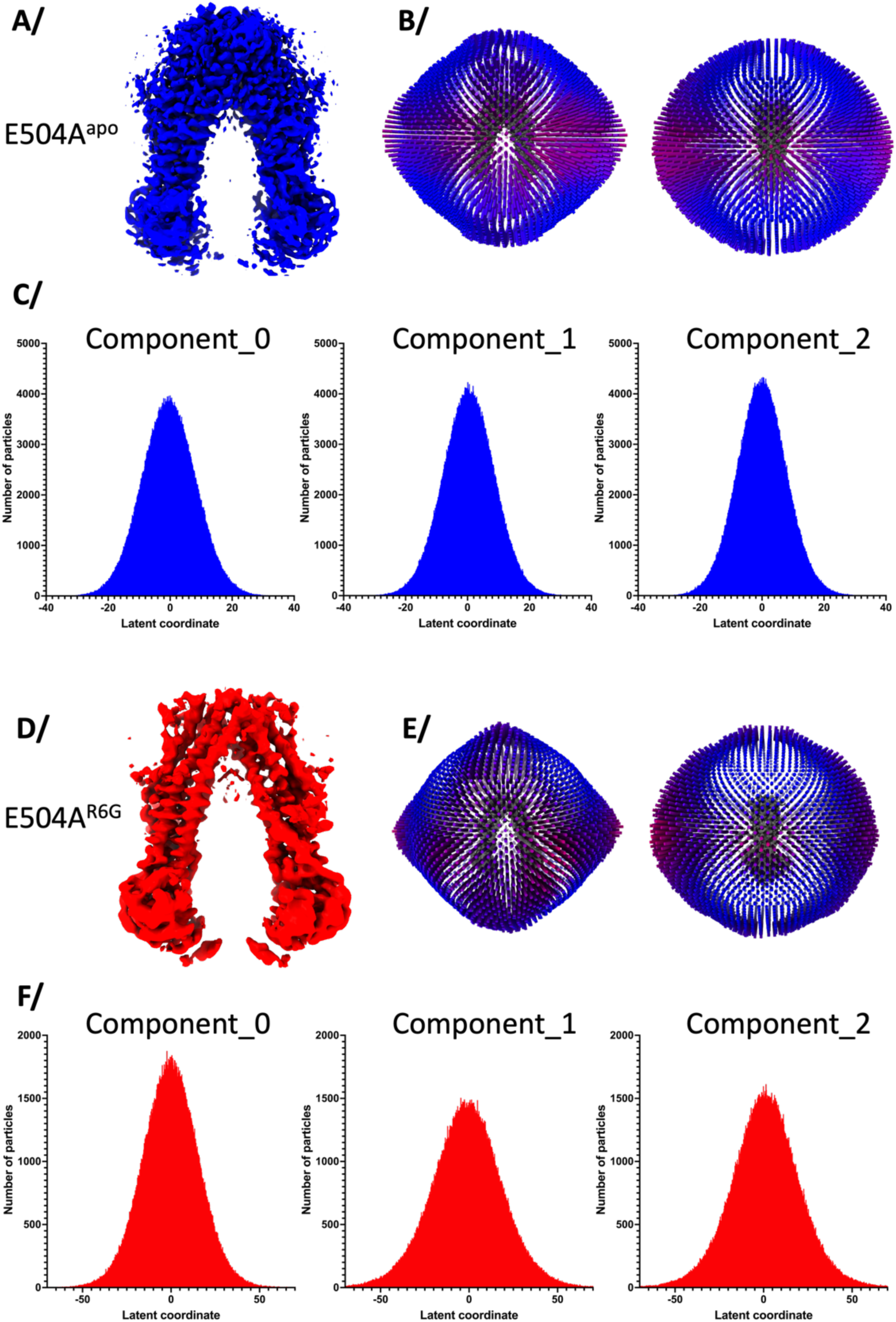
Particle distributions. **A/** E504^apo^ consensus reconstruction. **B/** 3D representation of particle distribution for the reconstruction; 2 views at 90° are represented. **C/** Particle distribution in latent space, represented per principal component as a function of latent coordinate. **D/ E/ F/** same as above for E504A^R6G^.

**Supp-Figure 13:**
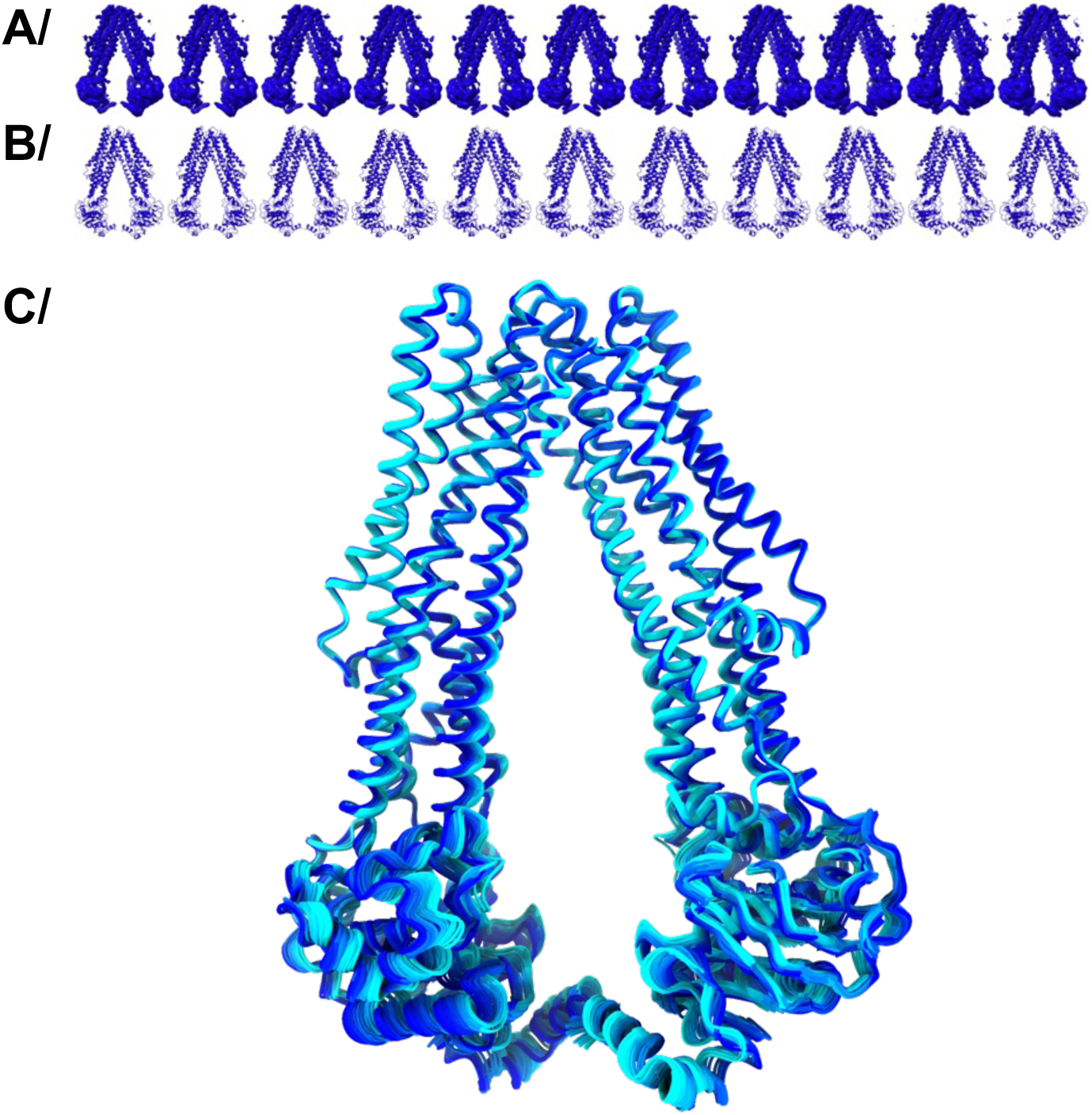
Explanation of *varref* output from 3DVA analysis. **A/** 11 evenly spaced out of the 20 maps output from 3DVA analysis in cryosparc for the condition BmrA^apo^. **B/** In each map, a model was refined by *phenix.varref*. The difference in models can be followed at the lowest part of the structure in C-terminal helices. **C/** All 20 models output from *varref* are shown and colored from blue to cyan to show the movement undergone by the protein.

**Supp-Figure 14:**
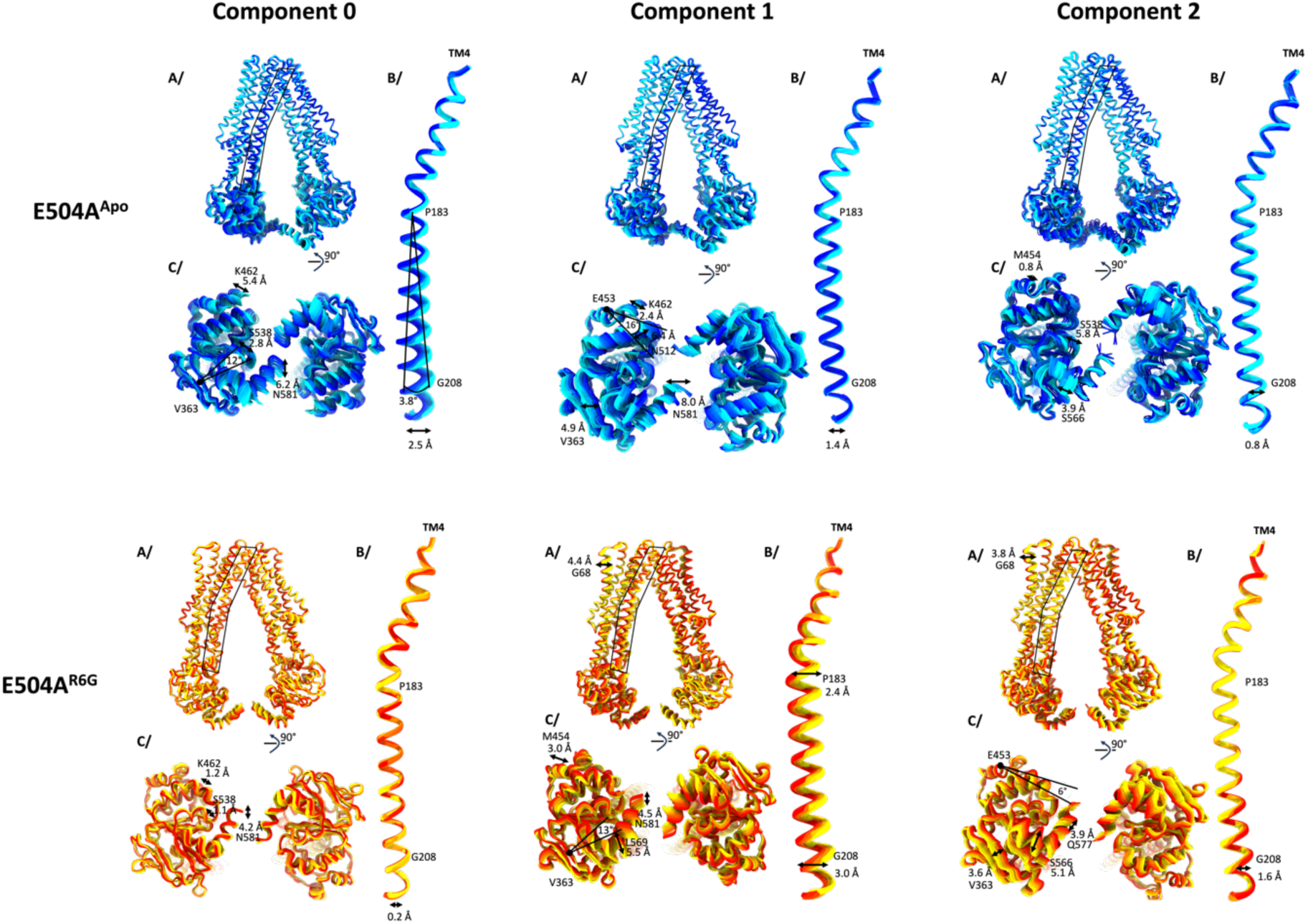
*varref* analysis for E504A^apo^ and E504A^R6G^. E504A^apo^ is represented in a gradient of blue to cyan and E504A^R6G^ from red to yellow. For each protein, the movement is decomposed in per principal component during the 3DVA analysis. For each panel **A/** is the lateral view of the full transporter, **B/** is the zoom on TM4, and **C/** is the view of the NBD from under. On each sub-panel, the rotation or translation is displayed and measured.

**Supp-Figure 15:**
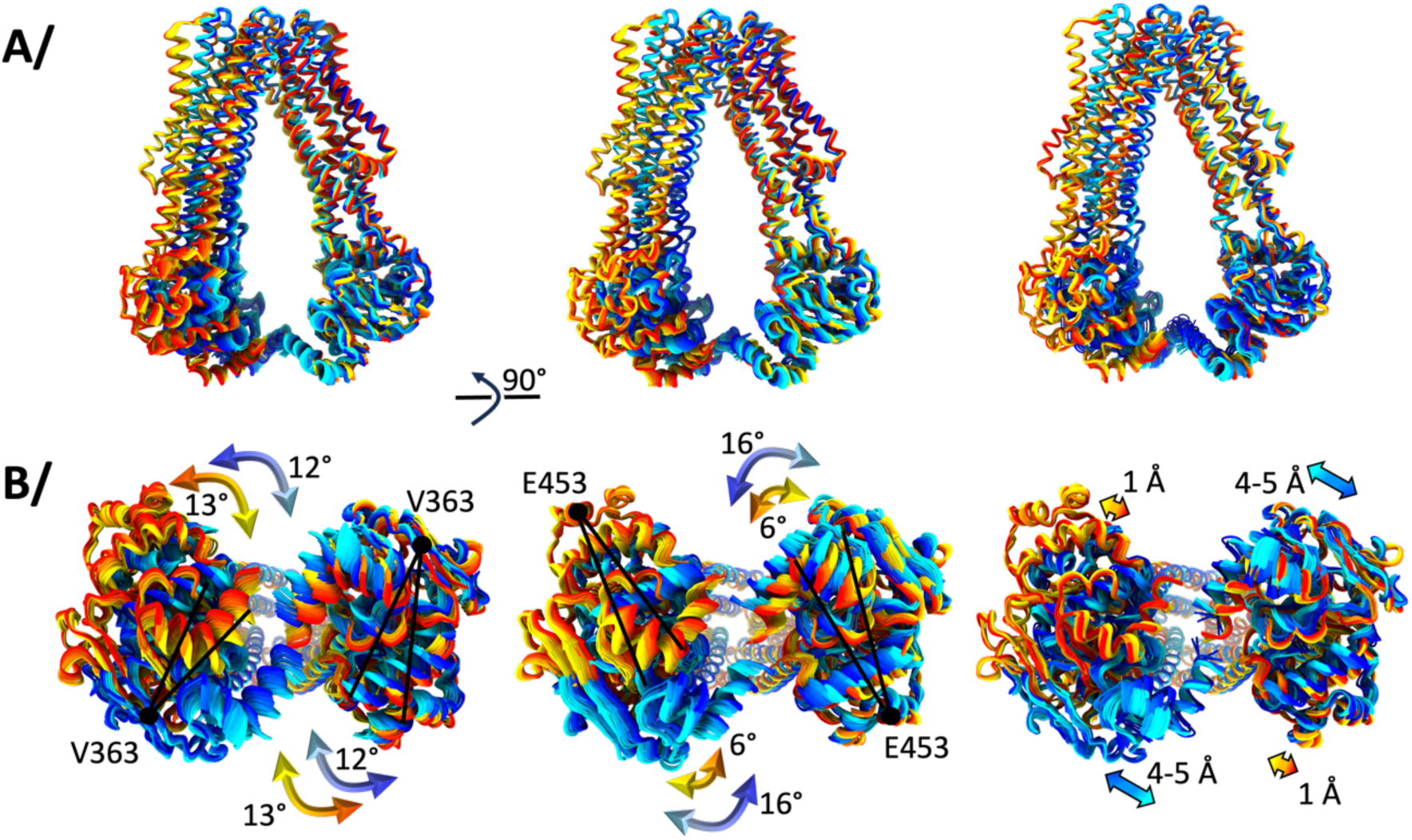
*varref* analysis for E504A^apo^ and E504A^R6G^. Same analysis as Supp-Figure 14 this time overlaid. **A/** lateral view and **B/** view from underneath. E504A^apo^ colored from blue to cyan and E504A^R6G^ colored from red to yellow.

**Supp-Figure 16:**
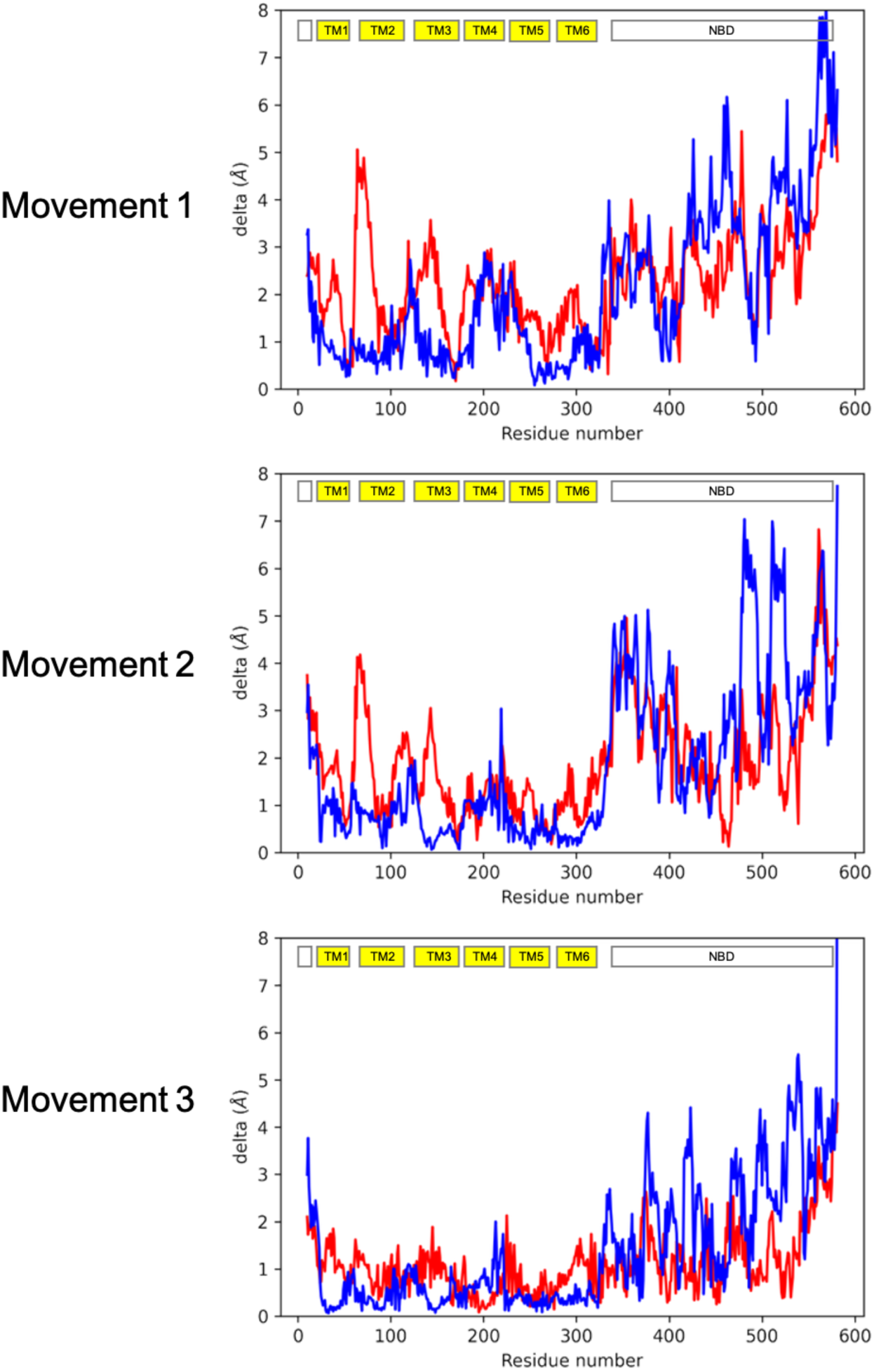
Structural variations within varref bundles. For each movement described in Figure 3A, rmsf was calculated for each structure within the bundle and plotted as a function of residue number in the chain. An average was performed between chain A and B of the homodimer. BmrA^apo^ is colored blue and BmrA^R6G^ is colored red. The position of each structural element is depicted on top of each graph for reference.

**Supp-Figure 17:**
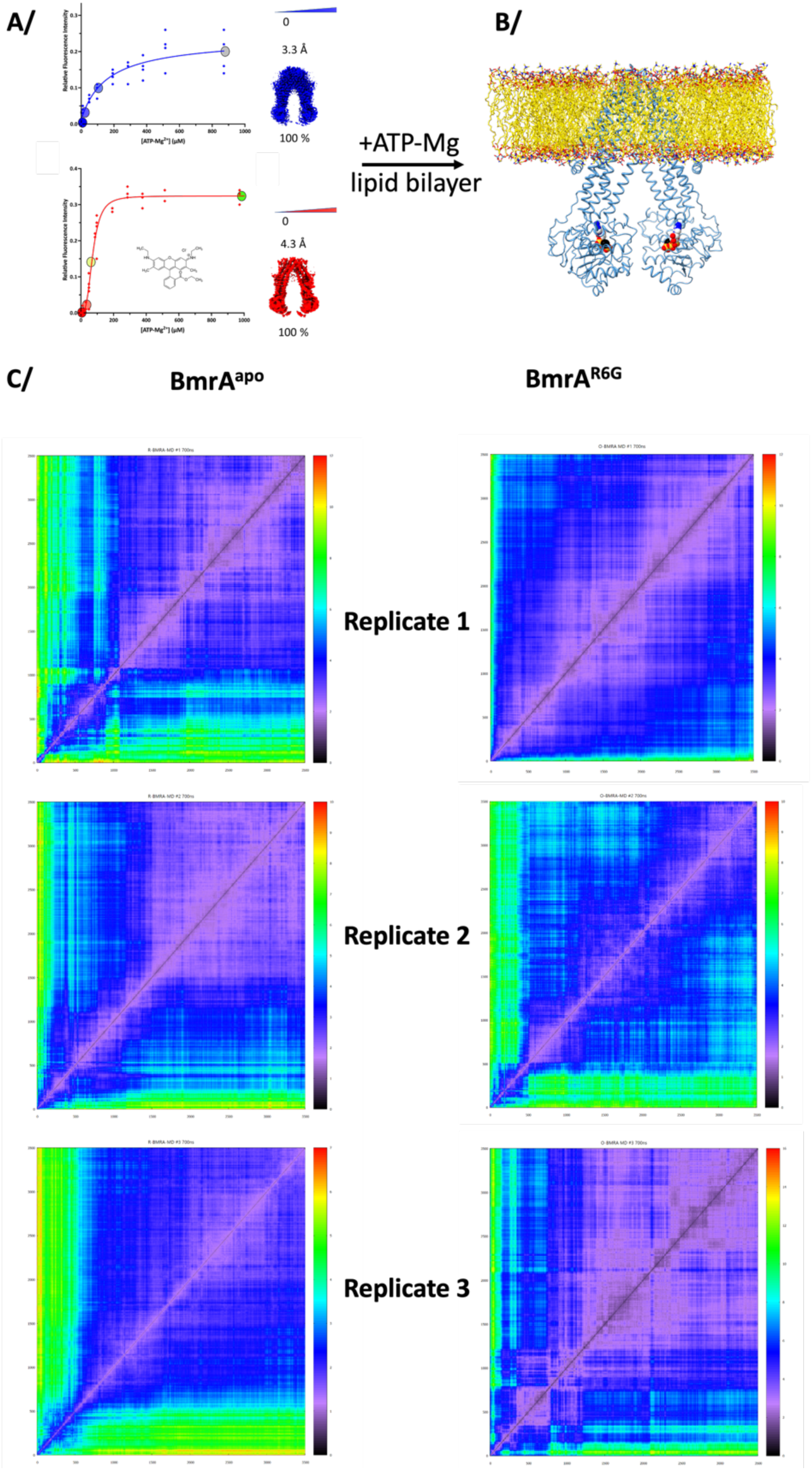
Global flexibility analysis of MD simulations. **A/** the two initial model used for MD simulations were the models obtained by cryoEM without ATP. **B/** ATP-Mg^2+^ was added in each binding site by superposition with the OF conformation in presence of ATP-Mg^2+^, and both proteins were inserted in a lipid bilayer and equilibrated before simulation. **C/** 2D rmsd plots across the 700ns long of simulations. Three replicates were made for each BmrA form BmrA^apo^ and BmrA^R6G^. X and Y axes are the frames. Color code on the right for the global rmsb between each frame, from low in black to high in red.

**Supp-Figure 18:**
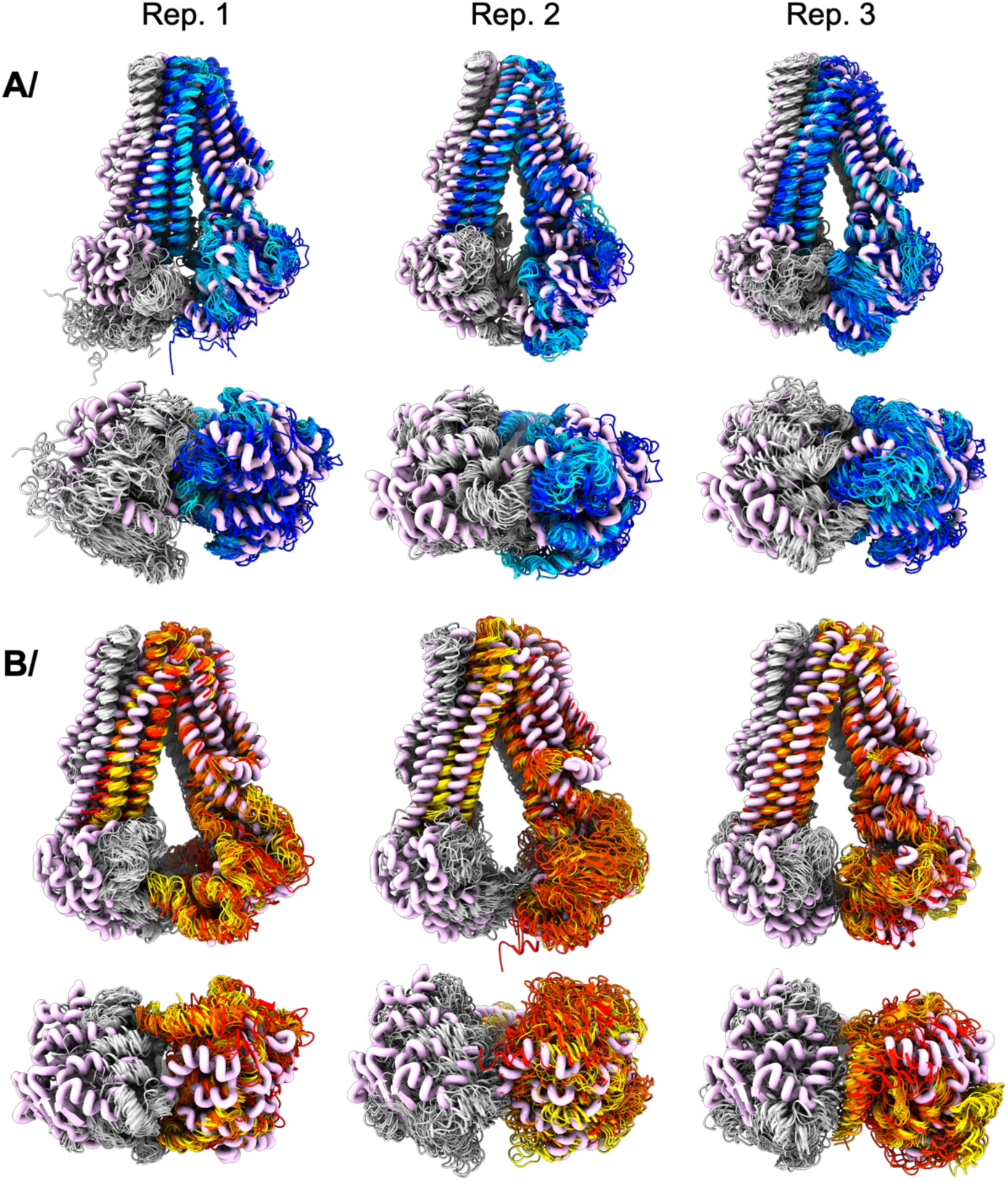
Visualization of protein movement during each MD simulation. **A/** E504A^apo^ was simulated in 3 independent replicas. The initial structure is shown as thick pink cartoon and a snapshot of the simulation is shown each 20 ns, colored from blue to cyan as the simulation progresses. **B/** Same as A/ but for BmrA^R6G^, colored from red to yellow.

**Supp-Figure 19:**
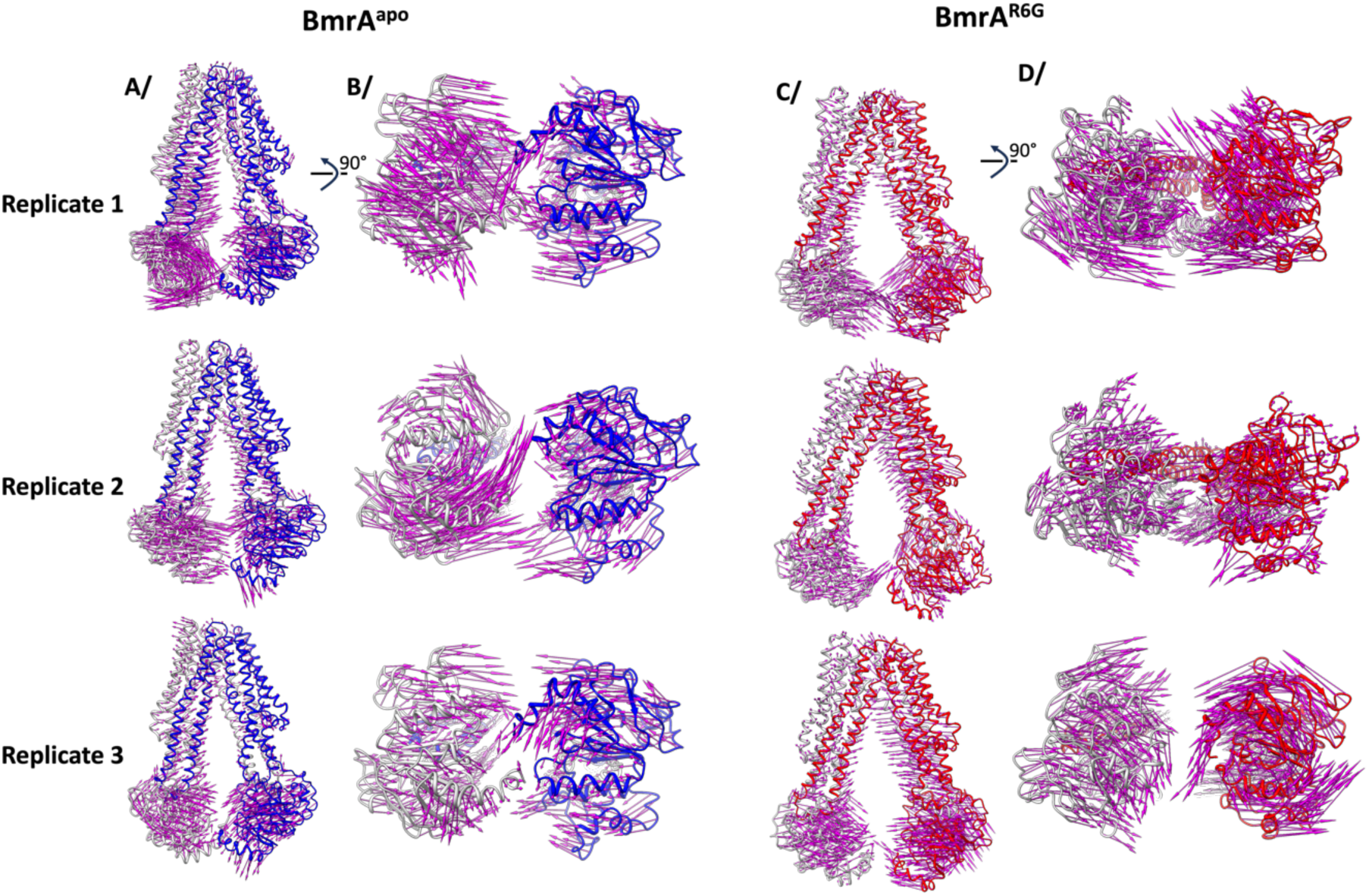
Visualization of protein movement during MD simulations. Same simulations as supp-Figure 16, with the arrow representing the distance between the initial and final frame of the simulation for E504A^apo^. One monomer is colored in blue, the other in light grey **A/** Seen from the side, and **B/** seen from under. **C/** and **D/** are the same as A/ and B/ for E504A^R6G^.

**Supp-Figure 20:**
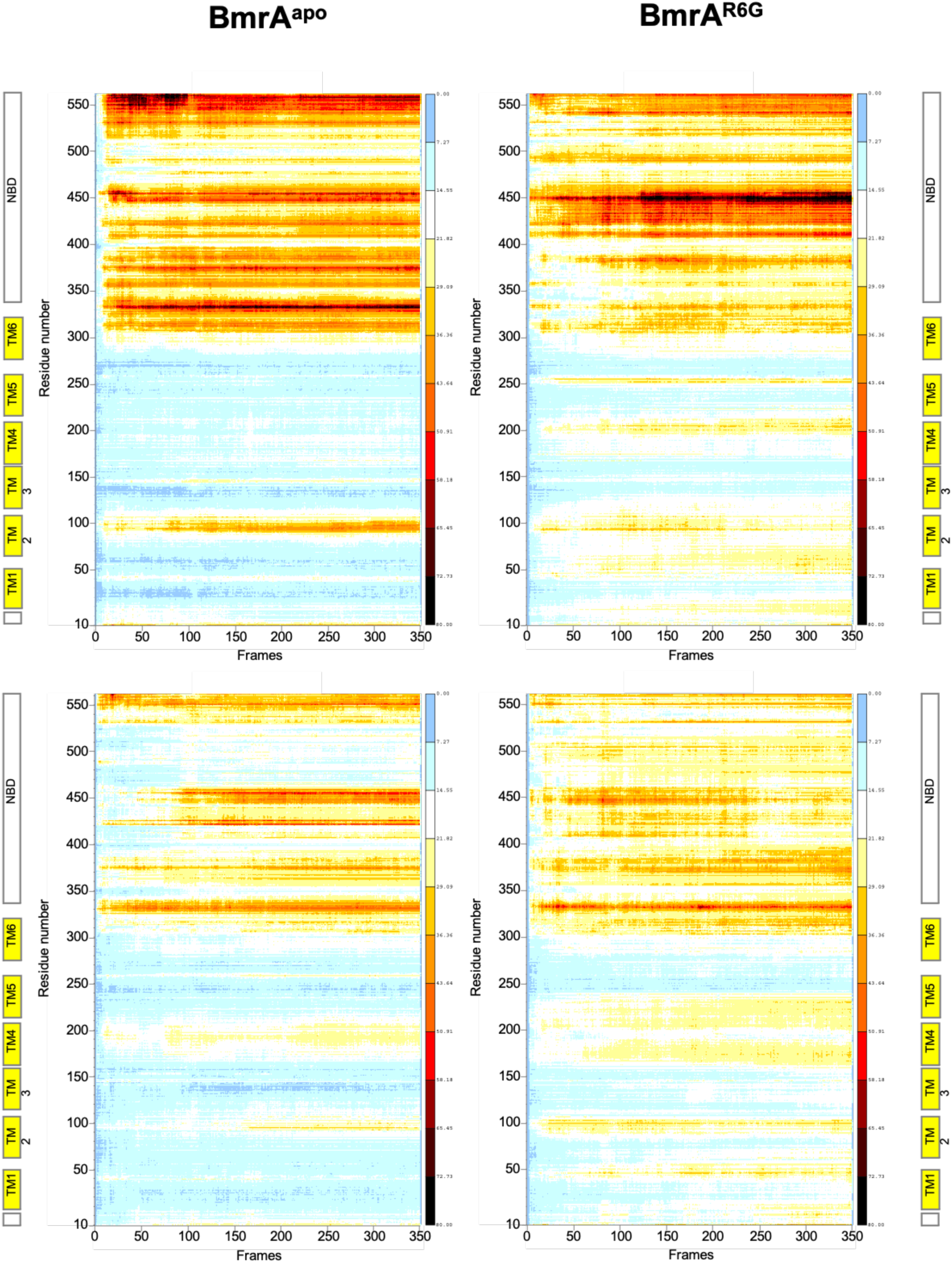
Residue displacement during MD simulations. For each protein E504A^apo^ or E504A^R6G^, the sum of displacement by residue across the three dynamics is displayed as a function of time. Each frame is sampled each 2ns, for a total of 700ns for each simulation. The color scale on the right represents the amount of displacement in Å, from 0 (blue) to 80 Å (black). The top and bottom panels represent the two halves of the transporter.

**Supp-Figure 21:**
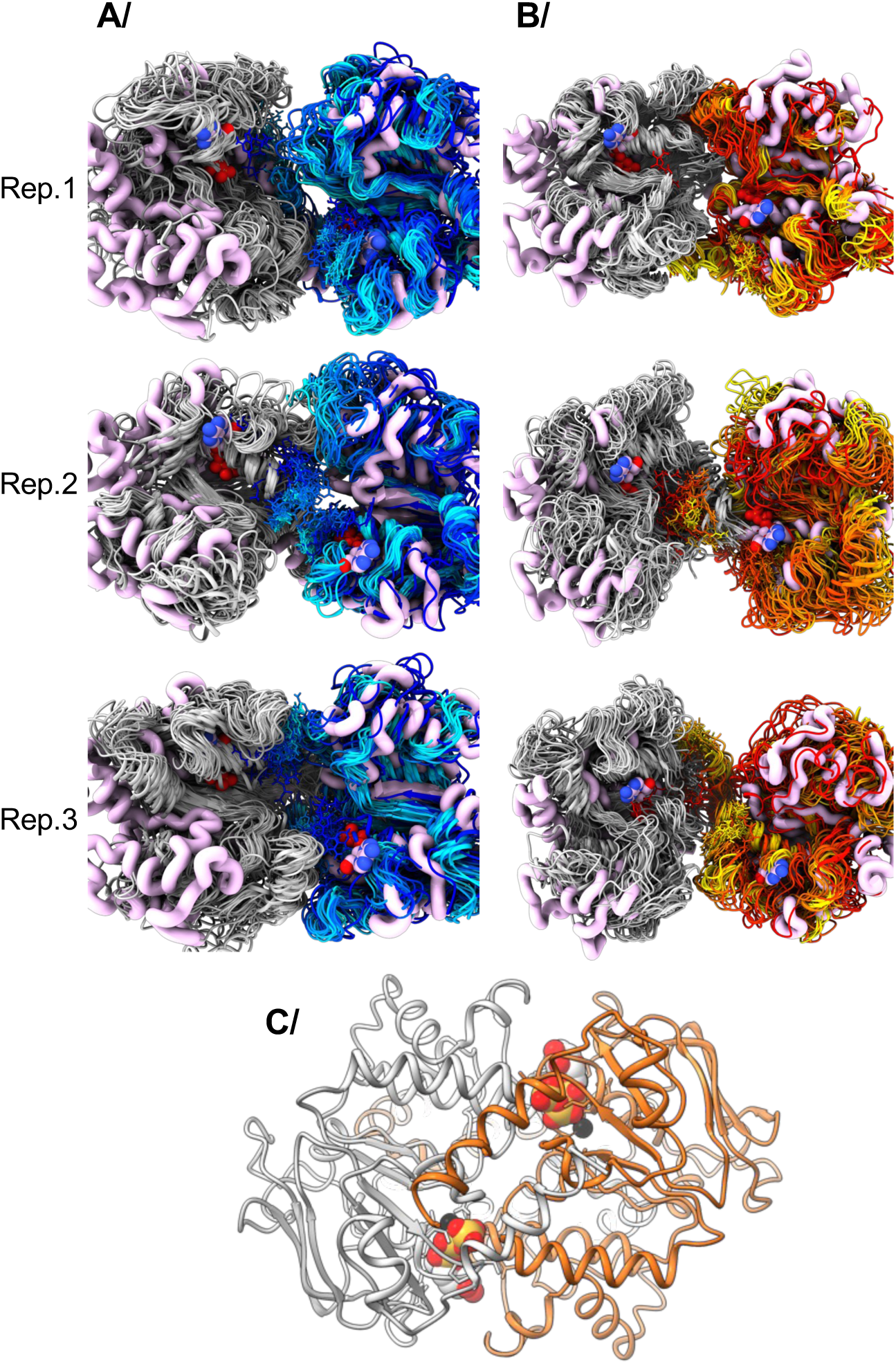
NBD and ATP flexibility during MD simulations. **A/** BmrA^apo^. **B/** BmrA^R6G^. For each replicate, the initial structure is shown in thick pink cartoon and initial ATP-Mg^2+^ position in thick sticks colored by atom type. The simulation of 700ns has been divided in 35 snapshots separated at equal time during the simulation, and represented in cartoon for the protein and sticks for ATP-Mg^2+^. The colors for the simulation range from blue to cyan or red to yellow to match the observations by 3DVA. **C/** OF conformation of BmrA for reference (PDB 7bg4), one monomer in orange, the other silver, ATP in spheres colored by atom type, and Mg in black.

**Supp-Figure 22:**
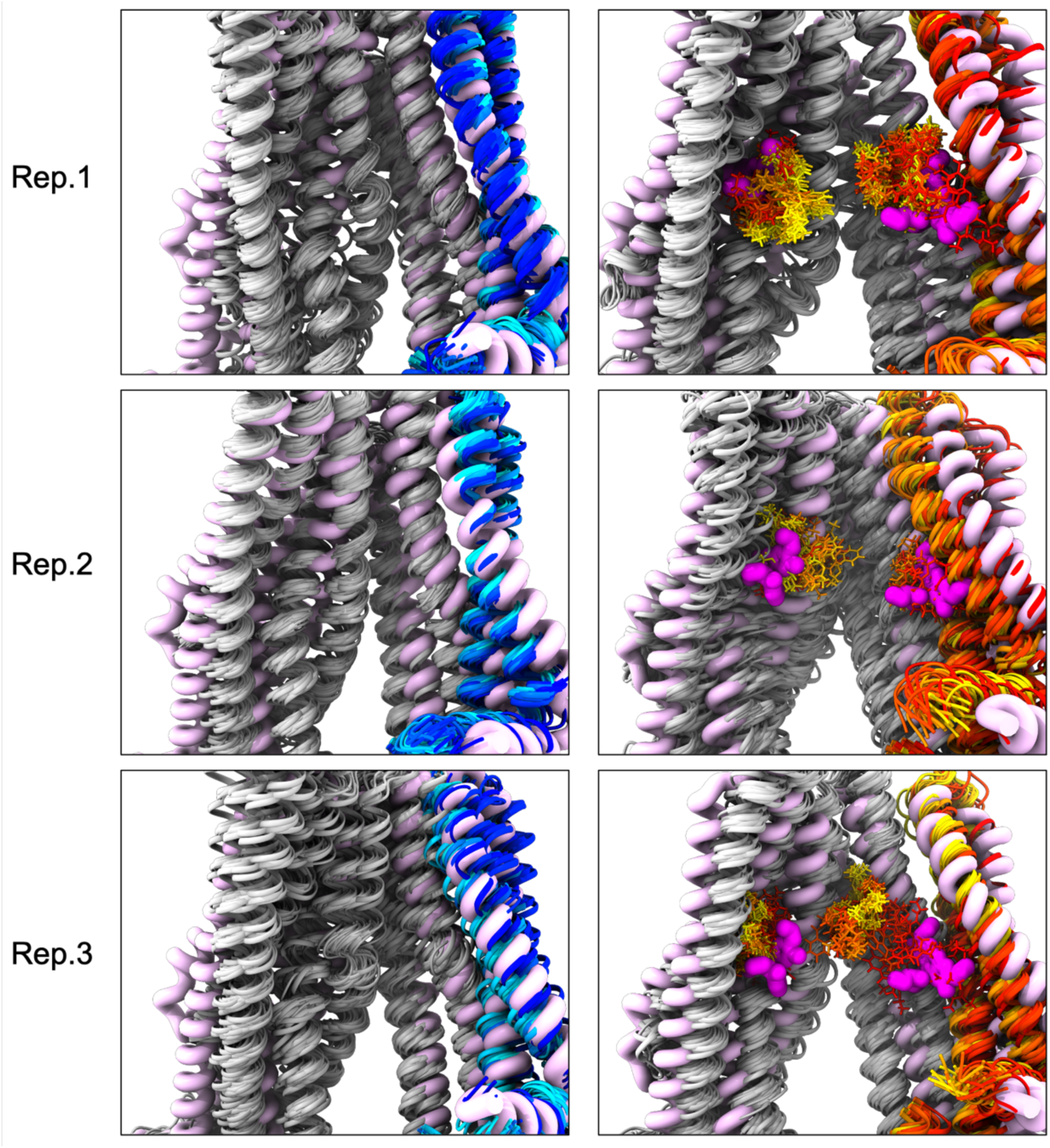
Mobility of R6G in its binding pocket. For each replicate of BmrA^R6G^, the initial structure is shown in thick pink cartoon and initial R6G position in magenta thick sticks. The simulation of 700ns has been divided in 35 snapshots separated at equal time during the simulation, and represented in cartoon for the protein and sticks for R6G. The colors for the simulation range from red to yellow to match the observations by 3DVA. The 3 replicates were made. Left panel, no R6G was present.

**Supp-Figure 23:**
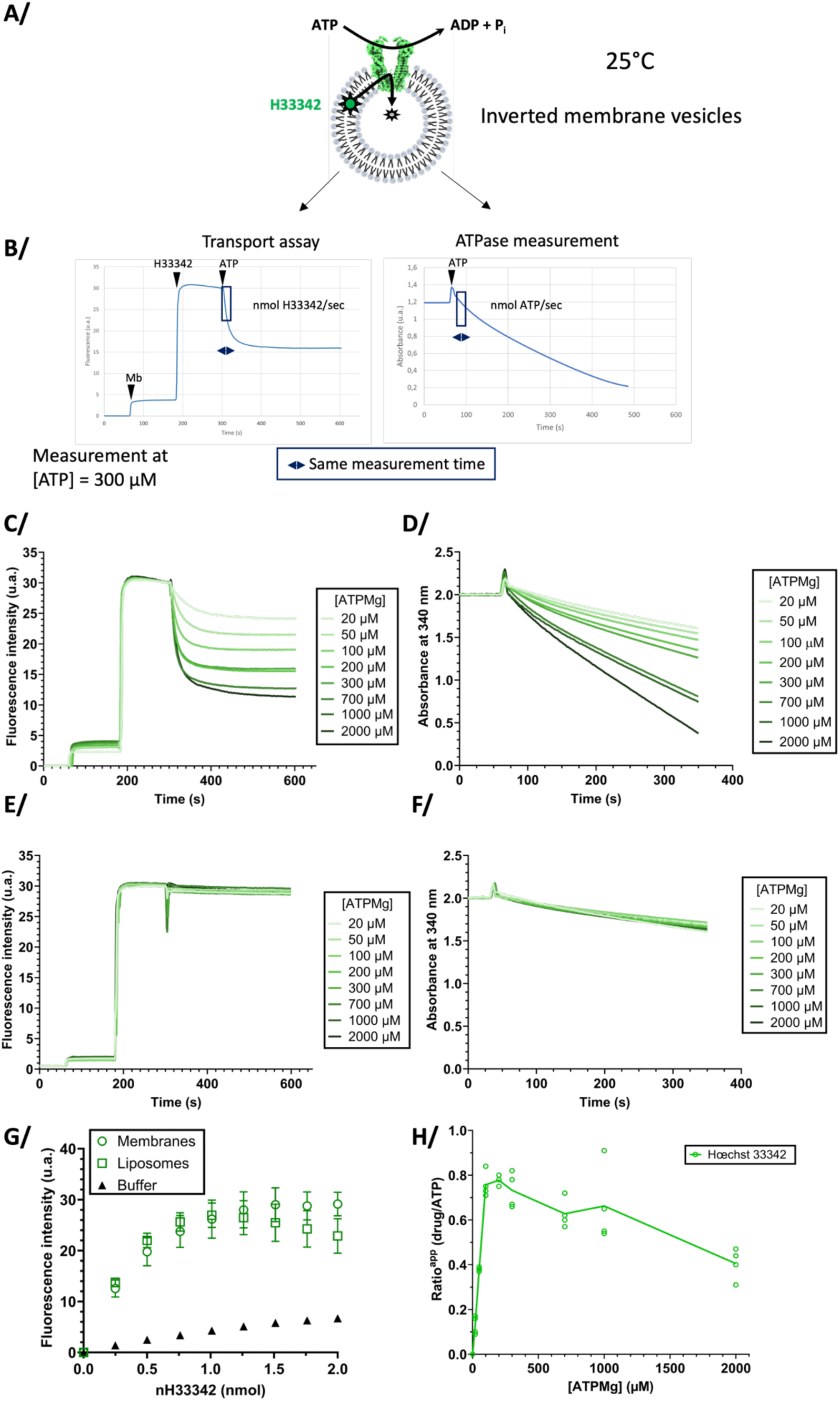
transport of Hœchst 33342 and ATPase activity for BmrA WT. **A/** Schematic representation of the experiment. **B/** Representative traces of substrate transport (left) and ATPase activity (right) for BmrA WT performed on the same membrane vesicles, the same day. All experiments have been carried out in quadruplicates, from 2 different membrane batches. The time range of measurement for transport is shown as a black rectangle on each graph. **C/** One monoplicate of substrate transport for the whole range of ATP tested. **D/** corresponding ATPase activities on the same membranes. **E/ F/** same as C/ and D/ but for the inactive mutant E504A. contaminant activities measured with the mutant E504A were deduced to BmrA WT activities for both transport and ATPase activities. **G/** Standard curve of Hoechst33342 fluorescence increase as a function of Hoechst33342 being added to a fixed amount of lipid, done on membranes (green circles), liposomes (green squares; liposome concentration chosen to have the same maximum as membranes) or in buffer (black diamond), for quantification of Hoechst 33342 transport. **H/** Ratio of Hoechst 33342 transported per ATP hydrolyzed, for each ATP concentration investigated.

**Supp-Figure 24:**
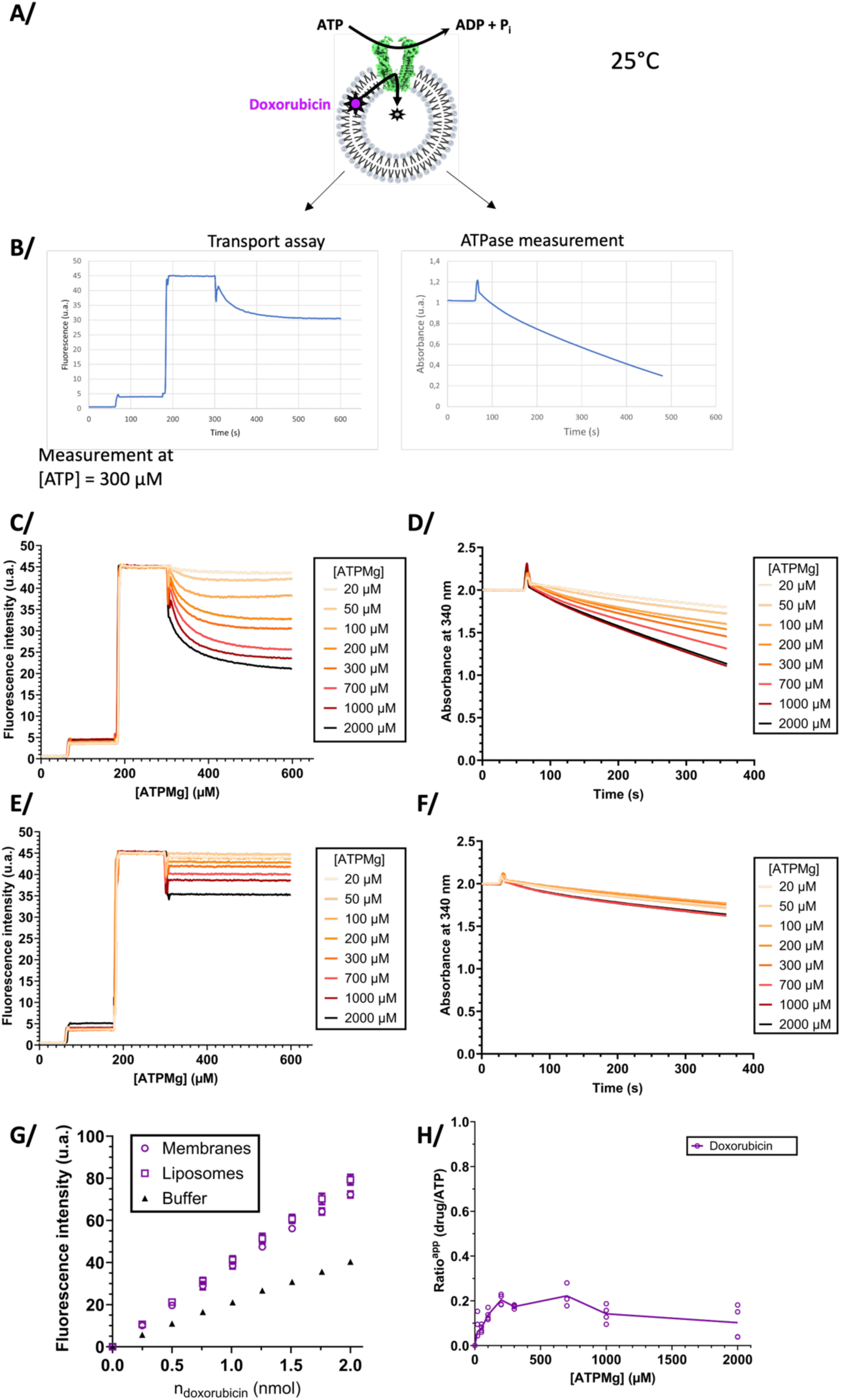
Doxorubicin transport. **A/** Schematic representation of the experiment. **B/** Representative traces of substrate transport (left) and ATPase activity (right) for BmrA WT performed on the same membrane vesicles, the same day. All experiments have been carried out in quadruplicates, from 2 different membrane batches. The time range of measurement for transport is shown as a black rectangle on each graph. **C/** One monoplicate of substrate transport for the whole range of ATP tested. **D/** corresponding ATPase activities on the same membranes. **E/ F/** same as C/ and D/ but for the inactive mutant E504A. contaminant activities measured with the mutant E504A were deduced to BmrA WT activities for both transport and ATPase activities. **G/** Standard curve of Doxorubicin fluorescence increase as a function of Doxorubicin being added to a fixed amount of lipid, done on membranes (green circles), liposomes (green squares, liposome concentration chosen to have the same maximum as membranes) or in buffer (black diamond), for quantification of Doxorubicin transport. **H/** Ratio of Doxorubicin transported per ATP hydrolyzed, for each ATP concentration investigated.

**Supp-Figure 25:**
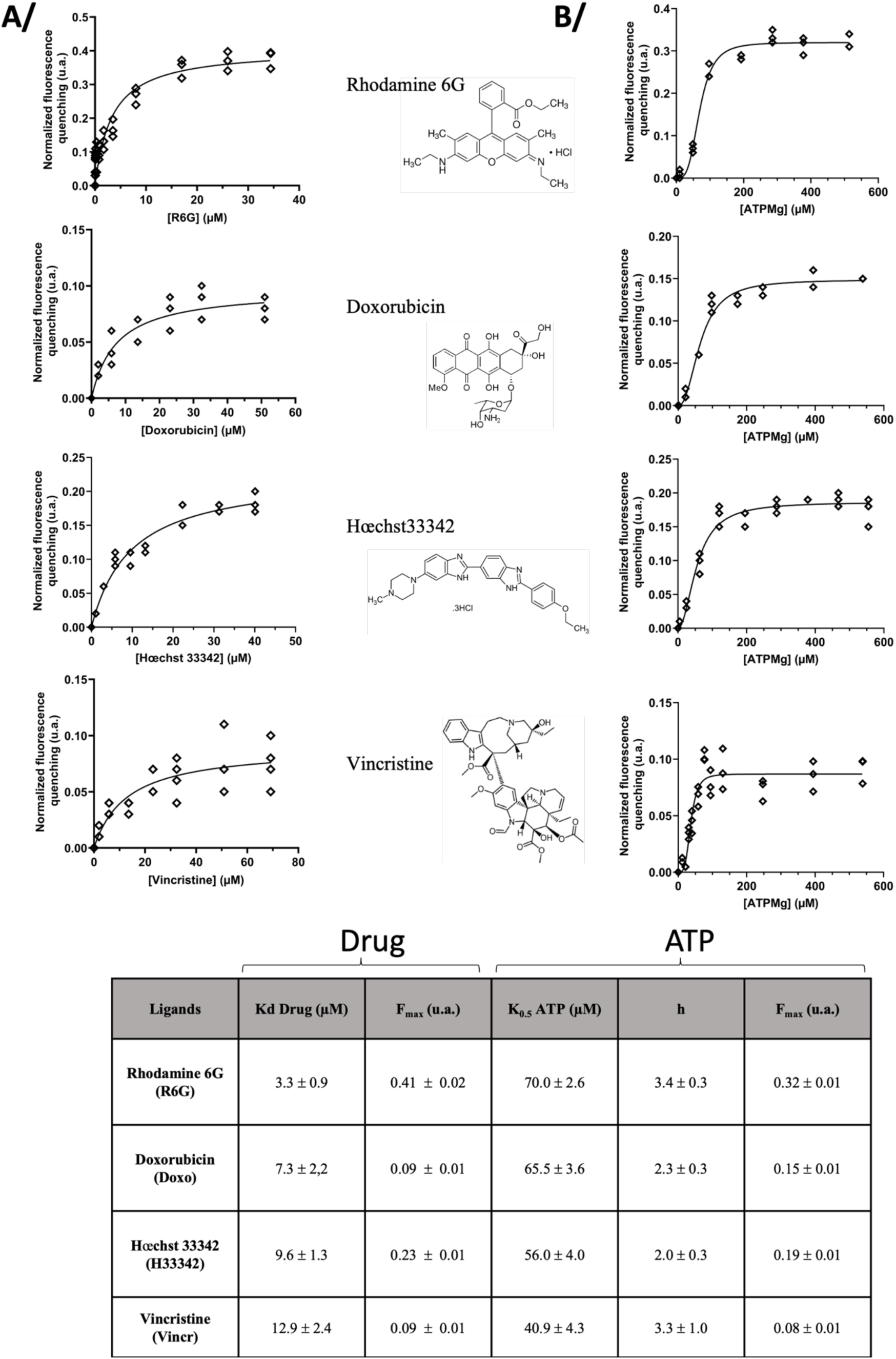
Drug and ATP binding on BmrA-E504A. Binding of R6G, Doxorubicin, Hœchst33342 or Vincristine were measured using intrinsic fluorescence displacement. Normalized fluorescence quenching is presented. B/ ATP-Mg^2+^ binding on BmrA-E504A in presence of drug, measured by intrinsic fluorescence, normalized quenching. The table recapitulates the Kd (drug), K_0.5_ (ATP), Hill number (h) and the maximum fluorescence quenching (F_max_).

